# Energy landscapes and heat capacity signatures for monomers and dimers of amyloid forming hexapeptides

**DOI:** 10.1101/2023.05.17.541223

**Authors:** Nicy, David J. Wales

**Affiliations:** Yusuf Hamied Department of Chemistry, University of Cambridge, Cambridge, CB2 1EW, U.K.

## Abstract

Amyloid formation is a hallmark of various neurodegenerative disorders. In this contribution, energy landscapes are explored for various hexapeptides that are known to form amyloids. Heat capacity (C*_V_*) analysis at low temperature for these hexapeptides reveals that the low energy structures contributing to the first heat capacity feature above a threshold temperature exhibit a variety of backbone conformations for amyloid forming monomers. The corresponding control sequences do not exhibit such structural polymorphism, as diagnosed via end-to-end distance and a dihedral angle defined for the monomer. A similar heat capacity analysis for dimer conformations obtained using basin-hopping global optimisation, shows clear features in end-to-end distance versus dihedral correlation plots, where amyloid-forming sequences exhibit a preference for larger end-to-end distances and larger positive dihedrals. These results hold for sequences taken from tau, amylin, insulin A chain, a de-novo designed peptide, and various control sequences. While there is a little overall correlation between the aggregation propensity and the temperature at which the low-temperature C*_V_* feature occurs, further analysis suggests that the amyloid forming sequences exhibit the key C*_V_* feature at a lower temperature compared to control sequences derived from the same protein.

## I. INTRODUCTION

Amyloid fibrils are associated with serious disorders, such as Alzheimer’s, Parkinson’s, type II diabetes and dialysis related amyloidosis. These amyloids are formed from specific proteins and the oligomers are toxic.^1^ The amyloid fibrils have a cross-beta structure that is formed by H-bonding between the NH and CO groups of the main chain of partially folded or misfolded proteins.^2, 3^ Since it is generally possible for proteins to establish such interactions between main chain atoms, amyloid formation has been suggested to be a generic property.^4^ However, the amino acid sequence also plays an important role, and amyloid formation propensity may be a sequence specific property.^5, 6^ Both amyloid forming proteins and short segments of these proteins can form steric zippers.^7–10^ All the peptides that form fibrils also form stable dimers.^11^ The aim of the present work is to investigate whether the propensity for amyloid formation is encoded in the energy landscape of monomers and dimers, and to provide a thermodynamic diagnostic involving the heat capacity (C*_V_*), which would complement models of amyloid formation based on the interactions within side chains.

The side chains of amino acid residues play an important role in amyloid formation,^12–14^ as they participate in aromatic–aromatic,^15^ electrostatic,^16^ and van der Waals interactions.^17, 18^ The hydrophobicity, secondary structure propensity, overall charge,^19^ exposed surface, dipole moment, and cooperativity in peptides correlate with amyloid formation ability.^20^ Deposition of peptides on amyloid templates is stereospecific and it involves interactions between peptide backbone and/or side chains.^21^ Several predictor algorithms have been developed to identify amyloidogenic proteins based on the insights obtained from amyloid structure and properties of amino acid residues.^22–25^ Different conformations of amyloid proteins can also occur at different temperatures and in different regions of the brain.^26^ These alternative conformations result in amyloid polymorphism.^27, 28^

However, the thermodynamic driving force for amyloid formation is still not well understood. Amyloid formation is studied mainly from the kinetic perspective.^29, 30^ Several attempts have been made to study the heat capacity of proteins^31–33^ using isothermal titration calorimetry and differential scanning calorimetry, and investigate the thermodynamics of amyloid formation.^34, 35^ Our recent heat capacity calculations have provided some insight into how the presence of tyrosine and arginine increases the phase separation propensity of proteins.^36^ This approach is also useful to rationalise the context-dependent properties of amino acids residues in different sequences.

Here, we explore the energy landscapes and calculate the heat capacity for monomers (Tables I and II) and dimers (Table I) of hexapeptides that are experimentally known to form amyloids, along with mutations that result in loss in amyloid formation ability (Fig. 1 and 2). We find that the low energy structures contributing to the first feature (peak or inflection point) above a threshold temperature in C*_V_* exhibit a variety of structures for the amyloid forming monomers (Fig. 3). Selected atoms in the hexapeptides are used to define end-to-end distance and dihedral value parameters. These parameters are useful to classify the variety of conformations that occur for the amyloid monomers (Fig. 4a). The low temperature feature in C*_V_* for amyloid monomers usually corresponds to a transition between structures with different backbone conformations with different main chain or side chain interactions. A similar heat capacity analysis for dimers shows another pattern in the end-to-end distance versus dihedral correlation plots. The low energy structures that contribute to the low temperature C*_V_* feature exhibit larger end-to-end distances and larger positive dihedrals for the amyloid forming sequences compared to the controls (Fig. 4b). We have also investigated the correlation between the aggregation propensity (calculated using Aggrescan^37^ software) and the temperature at which the first feature of interest occurs in the C*_V_* plot (Fig. 5). While there is little correlation between these two quantities, further analysis suggests that higher aggregation propensity correlates with lower temperature for the C*_V_* feature in sequences from the same protein. Extrinsic factors, such as pH, buffer conditions, and protein concentration^38^ are known to affect amyloid formation. The present study aims to complement these results by probing the thermodynamics of sequence specific properties for amyloid forming peptides. Water also contributes to interactions, but solvent dynamics are difficult to visualise and quantify,^39^ and here, water is modelled using an implicit solvent.

**FIG. 1:**
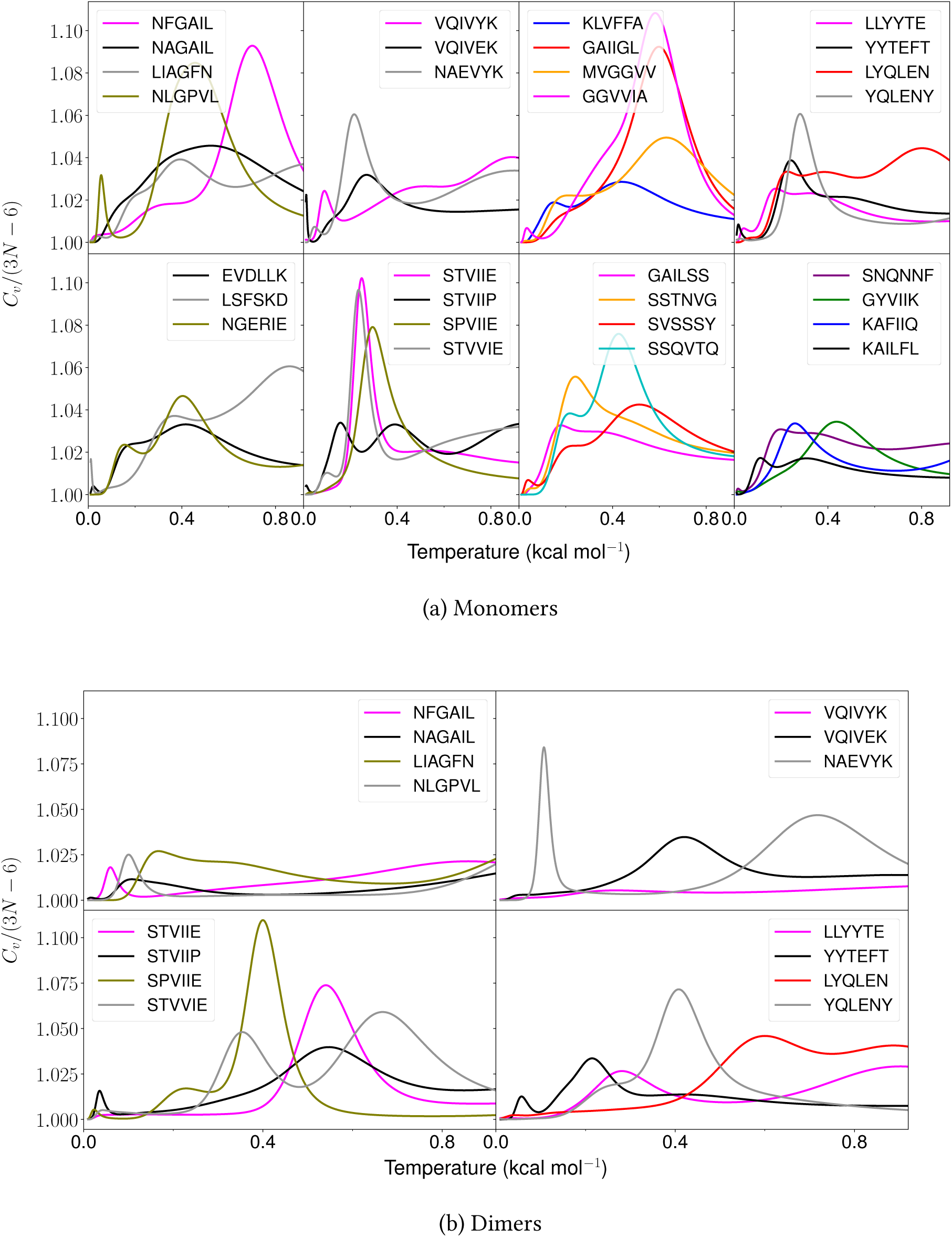
Heat capacity as a function of temperature (*k*_B_*T*) for various amyloid forming and control hexapeptide sequences.

**FIG. 2:**
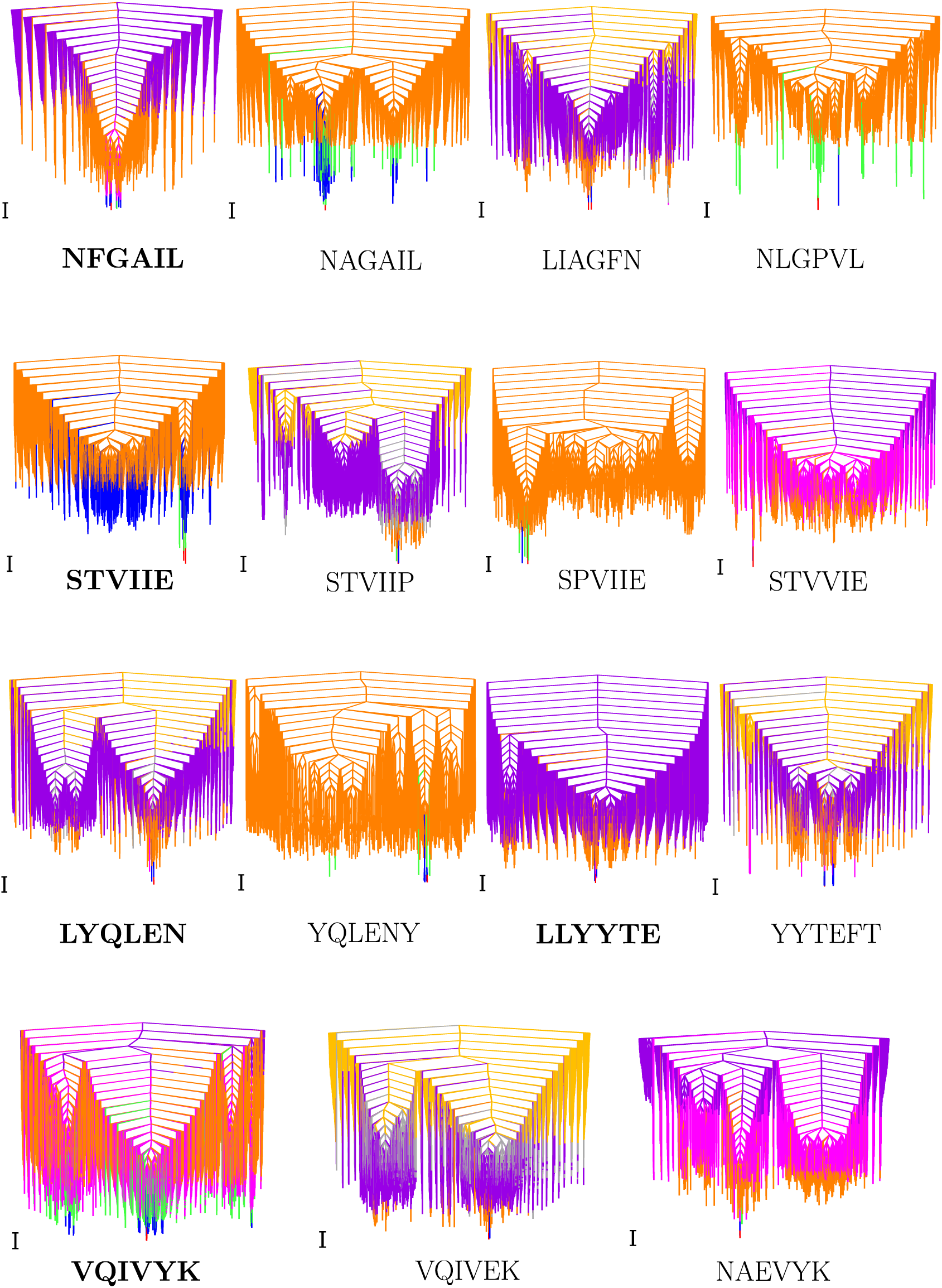
Disconnectivity graphs for amyloid (marked in bold) and control hexapeptide monomers. Local minima contributing to CV features (peaks/in.ection points) are represented by red to blue (feature 1), green to orange (feature 2), pink to purple (feature 3) and grey to yellow (feature 4). .e scalebar represents 1 kcal mol&100000;1.

**FIG. 3:**
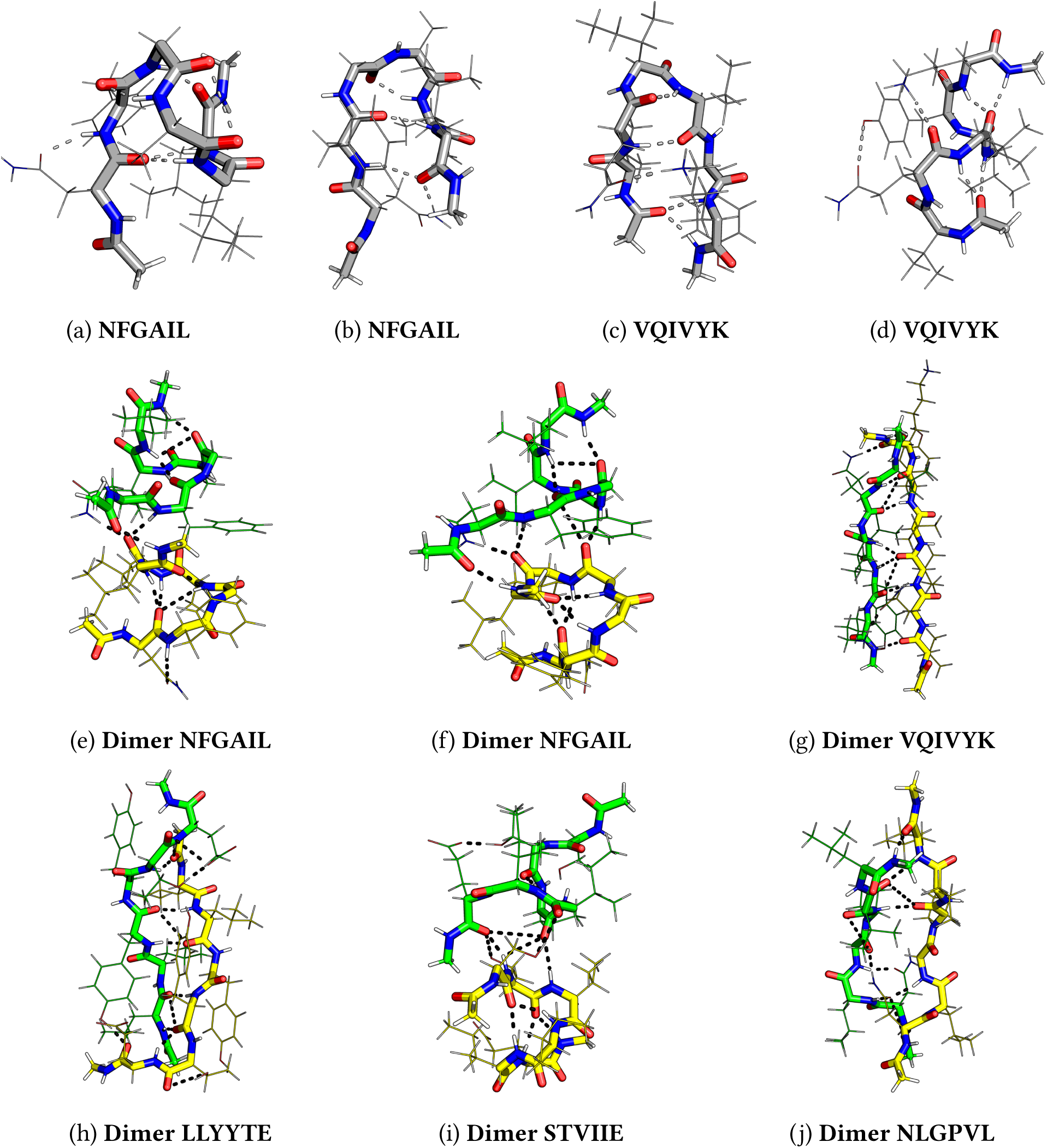
Various conformations of hexapeptide monomers and dimers that contribute to low temperature heat capacity feature.

**FIG. 4:**
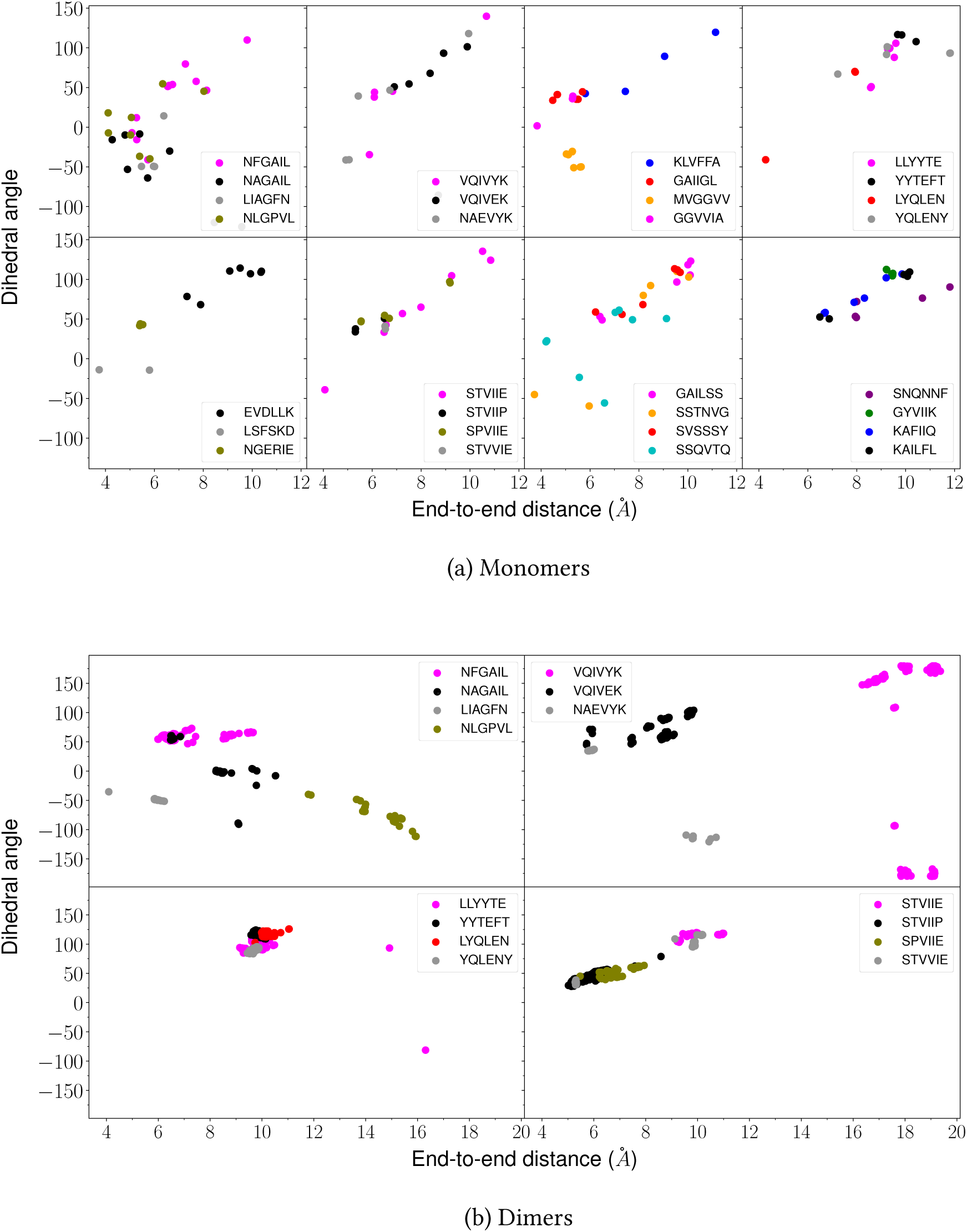
Correlation plot between end-to-end distance and dihedral angles for low energy structures contributing to first heat capacity peak of interest.

**FIG. 5:**
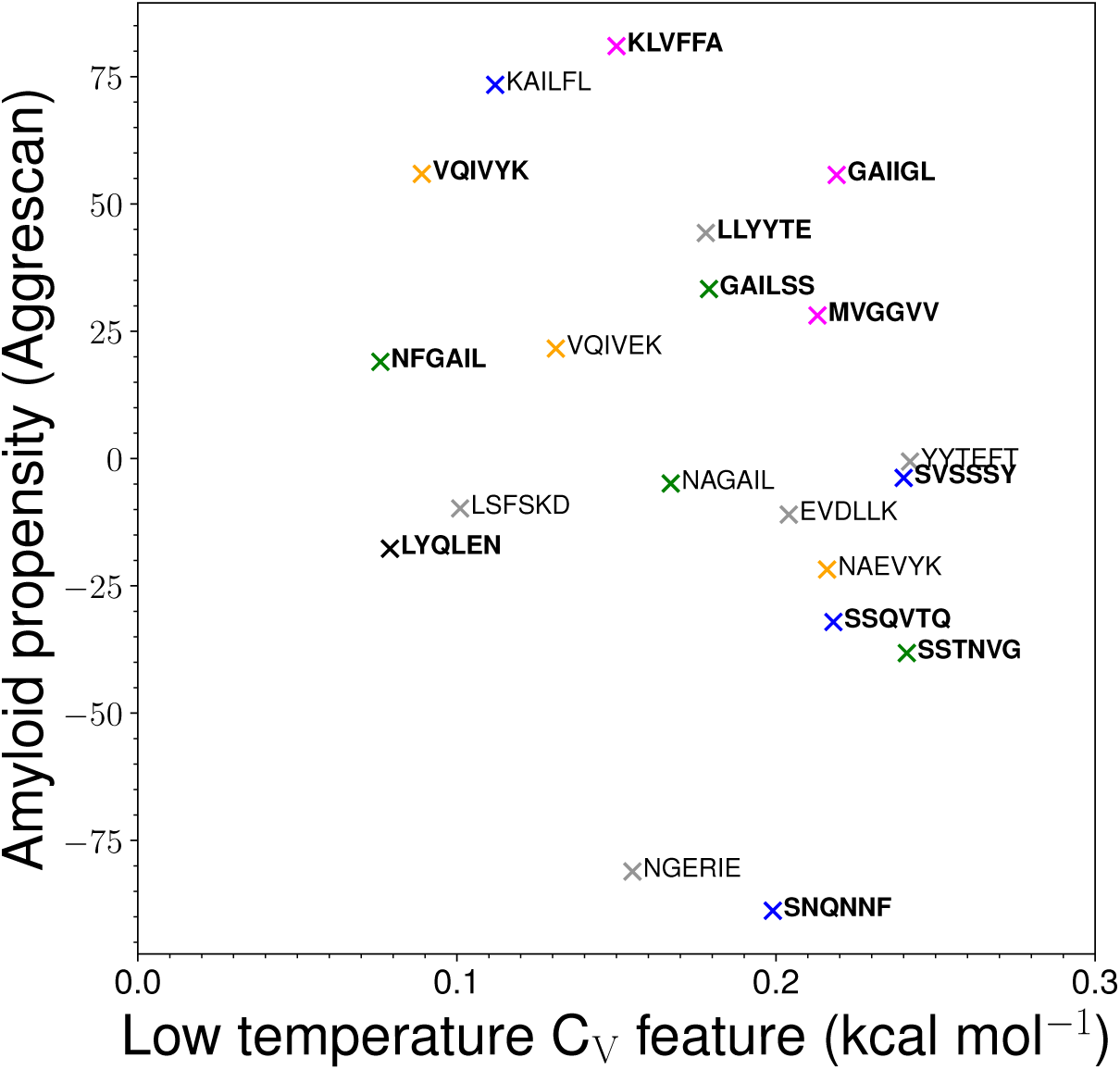
Correlation plot between the temperature (*k*_B_*T*) at which the first low temperature feature is observed (between 0.076–0.300 kcal mol*^−^*^1^) in the C*_V_* plot for the monomer and the propensity for amyloid formation predicted using the Aggrescan software^37^. The colours correspond to the proteins from which the sequences are taken. Orange, green, magenta, and grey correspond to tau, amylin, A-*β* and *β*_2_-microglobulin, respectively. Blue represents miscellaneous peptides taken from various proteins. Black represents a peptide which is amyloidogenic at a different pH. Several peptides are excluded from the above plot. The excluded peptides include GYVIIK, KAFIIQ, NLGPVL, and YQLENY, where the first C*_V_* feature is simply the melting peak; GGVVIA which has such a feature occurring above 0.3 kcal mol*^−^*^1^; the LIAGFN control sequence, for which the existing predictors fail to classify it differently from its reverse (NFGAIL) amyloid forming sequence; and the de-novo designed peptides STVIIE, STVIIP, SPVIIE, and STVVIE, which do not derive from naturally occurring amyloid forming proteins.

**TABLE I:**
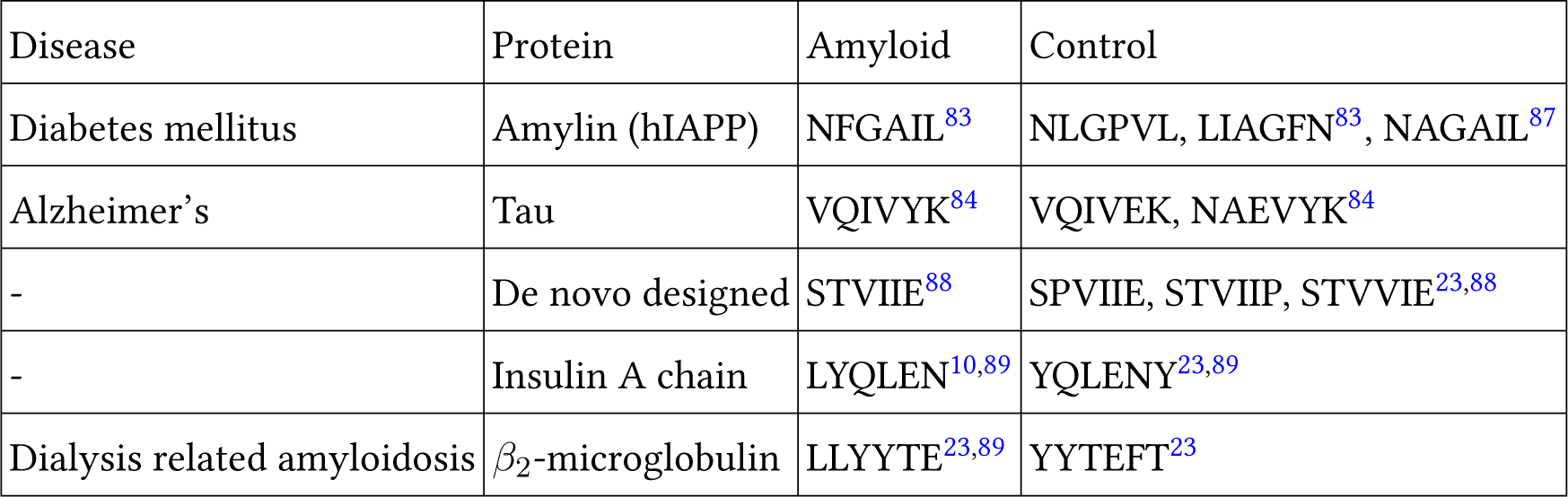
List of hexapeptides for which both the monomer and dimer energy landscapes have been explored.^10^,23,85,89–91

**TABLE II:**
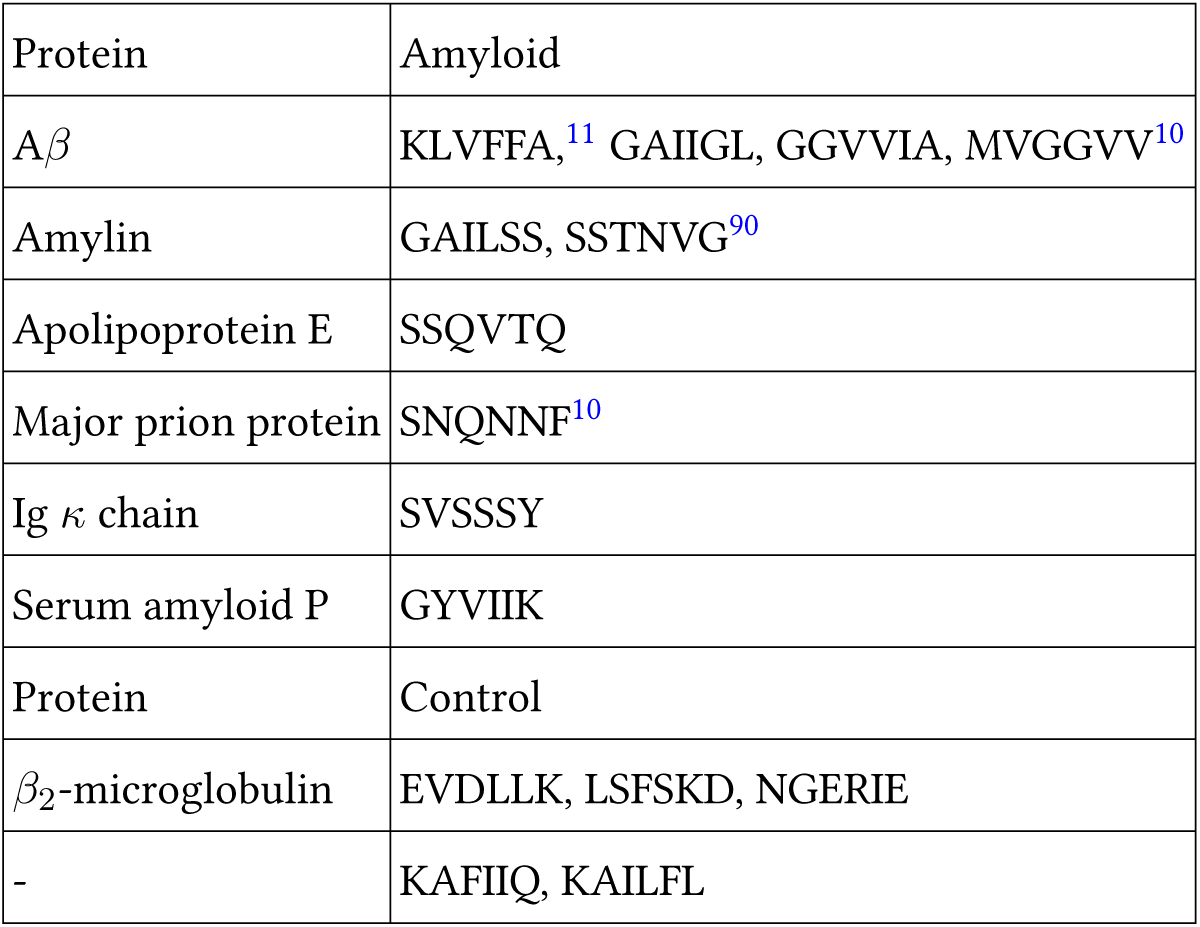
List of hexapeptides for which only the monomer energy landscapes have been explored.^10, 85, 90^

## **II.** METHODS

The hexapeptides are modelled using the FF99IDPs^40^ force field, which is a modified version of FF99SBILDN^41^ within AMBER.^42, 43^ This choice was made to gain better insight into the energy landscapes of various sequences using the same potential and compare it with our earlier study^36^ using the same force field. Our previous research^36^ showed that the structures that contribute to low temperature C*_V_* peaks for different AMBER force fields are similar. The N- and C-terminals of the peptide are methylated and methylamidated, respectively. The topology file is symmetrised^44^ to account correctly for permutational isomers. Water is modelled using implicit solvation (igb=8) along with a 0.1 M monovalent ion concentration.^42, 43^

The initial landscape exploration is performed using basin-hopping parallel-tempering (BHPT)^45–47^ implemented within the global optimisation program GMIN^48^ interfaced with AMBER. Similar to our previous study^36^ on monomers, 16 replicas are used for BHPT, with temperatures exponentially distributed between 300–575 K. For dimers the potential energy landscapes are explored using a combination of rigid body, Cartesian, and group rotation moves.^49–51^ Each monomer is used to define a rigid body. The two monomers are expanded radially from the centre of coordinates, rotated within the angle-axis framework,^52^ and translated, with rigid body moves performed after every 111 Cartesian moves. The thousand lowest energy structures are saved and converged tighter to a root-mean-square convergence criterion for the gradient of 10*^−^*^7^ kcal mol*^−^*^1^ Å *^−^*^1^. A total of 600000–800000 basin-hopping steps are performed for each peptide dimer. This total involves different runs with different starting structures, step size, and frequency of rigid body moves. For dimers, the sampling was monitored using the convergence of low temperature heat capacity peaks.

For monomers, discrete path sampling^53^ was employed to obtain pathways for interconversion between local minima and the global minimum. Each multistep pathway is composed of minimum– transition state–minimum triples.^54, 55^ Several geometry optimisation algorithms are employed to obtain these pathways,^56^ including the doubly-nudged^57, 58^ elastic band^59–62^ algorithm, hybrid eigenvector-following,^63–66^ minimisation with a limited memory Broyden-Fletcher-Goldfarb-Shanno algorithm,^67, 68^ and Dijkstra’s shortest path algorithm.^69^ These tools are implemented within the OPTIM^70^ program. The PATHSAMPLE^71^ program is used to expand the stationary point database, including a strategy to remove unphysical barriers connecting various local minima.^72^ The convergence of the database is monitored using disconnectivity graphs and the convergence of low temperature heat capacity peaks.

Disconnectivity graphs^73, 74^ provide an overview of the landscape organisation. These graphs preserve the information about the highest energy barrier that needs to be overcome to interconvert a pair of connected local minima. The vertical axis represents the potential or free energy, and the nodes along the vertical axis define superbasins. The minima within a superbasin can interconvert by overcoming a barrier that is less than or equal to the threshold energy associated with the corresponding node. The branches terminate at the potential or free energy of individual local minimum.

The heat capacity of the peptides is estimated using the harmonic superposition approximation.^75–80^ The partition function of each local minima is obtained using normal mode analysis and the total partition function is the sum of partition functions of all the local minima. Each peak in the heat capacity involves contributions from minima with negative and positive occupation probability derivatives with respect to temperature.^81^ The principal contributions for each heat capacity peak are visualised in disconnectivity graphs using different coloured branches for each set of minima (Supplementary Information). The geometric parameters (end-to-end distance and dihedral angle) used to distinguish different structures are calculated using the CPPTRAJ program within AMBER.^82^

## **III.** RESULTS AND DISCUSSION

Various hexapeptides from naturally occurring proteins are found to be important contributors of protein aggregation, and these segments can themselves also form amyloids. For example, NFGAIL^83^ and VQIVYK^84^ occurring in human islet amyloid polypeptide (hIAPP or amylin) and tau protein, respectively, form amyloids, and the aggregates of these proteins are found in type II diabetes and Alzheimer’s, respectively. Here, we explore the potential energy landscapes of amyloid forming hexapeptides and control sequences to investigate if there is any incipient signature for this behaviour in the heat capacity, which tells us about the competition between alternative low-lying conformations that differ significantly in their enthalpy and entropy. Both the control and amyloid forming hexapeptides were found to exhibit features at low temperatures. Interestingly, the structures with varying backbone conformations were found to contribute significantly to the low-temperature feature in amyloidogenic peptides. Note that the feature (peak/inflection point) of interest is taken as the first feature that occurs above *k*_B_*T* = 0.086 kcal mol*^−^*^1^ in C*_V_* plot. This threshold is employed because the presence of specific residues (isoleucine, valine, leucine and tyrosine) leads to low-lying structures separated by relatively small barriers. These barriers occur due to the presence of different conformations for the side chains and different rotamers for tyrosine. The N-terminal and C-terminal caps and the residues at the ends of the peptides are relatively free to orient in different directions and establish H-bonding within the peptide in different ways. Such structures are also separated by relatively small barriers. Although these structures do not differ in the main chain conformations, they may give rise to features (peaks or inflection points) below *k*_B_*T* = 0.086 kcal mol*^−^*^1^ in C*_V_* . Hence, the low temperature peak of interest that represents a transition between structures with different main chain conformations is the first peak that occurs above the threshold temperature. The heat capacity analysis is performed on the converged landscapes (Fig. 1).

The energy landscapes for monomers, visualised using disconnectivity graphs,^73, 74^ can be multifunneled for both amyloid forming and control hexapeptides (Fig. 2). However, the structures lying at the bottom of funnels separated by significant barriers differ significantly for the amyloids and controls (Supplementary Information). The low energy minima separated by large barriers interconvert via breaking of H-bonds between main chain and side chain atoms, opening of the peptide backbone, and refolding, which results in a different backbone conformation.

Various conformations observed for hexapeptides appear similar to beta-hairpin (U-shaped), ?-shaped, S-shaped (partially helical), W-shaped (almost helical), and extended Z-shaped structures, which are listed in the order of increasing end-to-end distance (Fig. 3). The end-to-end distance is defined between the N atom of the first residue and the C atom of the sixth residue in a hexapeptide. The dihedral angle is defined between the C*_α_* atoms of the first, third, fourth and sixth residue in the hexapeptide. Small distance and small dihedral angle parameters correspond to hairpin-like structures, and large distance and large dihedral angle correspond to helical structures. Even larger distances signify extended monomer structures. These distance and dihedral angle parameters are calculated for the structures contributing to the low-temperature C*_V_* feature and we examine correlations with the amyloid formation propensity. The quantitative correlation between the exact temperature at which the C*_V_* feature occurs and aggregation is also investigated.

### Monomer heat capacity

Interesting patterns are found on analysing the funnels that contain structures contributing to the first C*_V_* feature (peak or inflection point) above *k*_B_*T* = 0.086 kcal mol*^−^*^1^ (Fig. 1a). In general, for amyloid-forming hexapeptide sequences, such as NF-GAIL, VQIVYK, STVIIE, LYQLEN and KLVFFA, several significantly different main-chain conformations occur in different funnels (Fig. 3). Both the helical and hairpin structures consistently appear together in the landscapes for amyloid hexapeptides. In contrast, for the controls (NAGAIL, VQIVEK, NAEVYK, YYTEFT, EVDLLK, LSFSKD, NGERIE, STVIIP, STVVIE, YQLENY, KAFIIQ, and KAILFL) only one (or a set of states with similar secondary structure) main chain conformation occurs at the funnel bottom. This result is also evident from the greater spread of points for amyloid forming sequences compared to the respective control sequences in the end-to-end distance versus dihedral correlation plot (Fig. 4a). Different side chain interactions may be present in the helical and hairpin structures of amyloids. Helical structures generally have a H-bond between side chains of two residues, while beta-hairpins have a H-bond between side chains of another pair of residues, with one participating residue being common in establishing two different kinds of H-bond patterns in the two different main chain conformations. This situation occurs for VQIVYK, STVIIE, and LYQLEN amyloid hexapeptides. The control sequences lack such interactions between side chains. It is the residues containing hydroxyl or amide groups in their side chains that usually participate in these key interactions.

However, there are some amyloid forming sequences that do not show a variety of low energy backbone conformations contributing to the first C*_V_* peak of interest. This situation occurs for GAILSS, SNQNNF (almost helical), MVGGVV (hairpin), SVSSSY, and GYVIIK (partial helical). GAIIGL and GGVVIA exhibit partial helical structures. LLYYTE does not form a proper hairpin that can be classified in terms of a small end-to-end distance. These observations are also evident from the narrower range of points for these peptides in the distance versus dihedral correlation plot (Fig. 4a). Similarly, the control sequences NAEVYK and SPVIIE, which at first do not appear to follow the trend shown by other control sequences, with a wider spread of points in the correlation plot, are found to follow the trend when their disconnectivity graphs are analysed. Both sequences lack contributions from helical structures. However, the occurrence of extended structures leads to the points with larger distance in the correlation plot. For SPVIIE, the peak of interest is the same as the melting peak and hence extended conformations are bound to occur at this temperature. In other words, SPVIIE does not show a low temperature peak, it shows just a melting peak.

Various intramolecular interactions occur between amino acid side chains and the backbone in the low-energy structures that contribute to features in C*_V_* . The residues with an -OH group, such as serine, threonine, and tyrosine, can establish H-bonds and interact via H atom with the -CO group (main chain) or -COO*^−^* group (aspartic or glutamic acid), and via the O atom with the -NH group (main chain or lysine). Similarly, the presence of an amide group in asparagine and glutamine allows the same residue to establish two different types of H-bonds, i.e., one in which the -CO group interacts and another in which the -NH group interacts. The specific interactions found between amino acid pairs in some of the low energy structures for the hexapeptide monomers are summarised in Table III.

**TABLE III:**
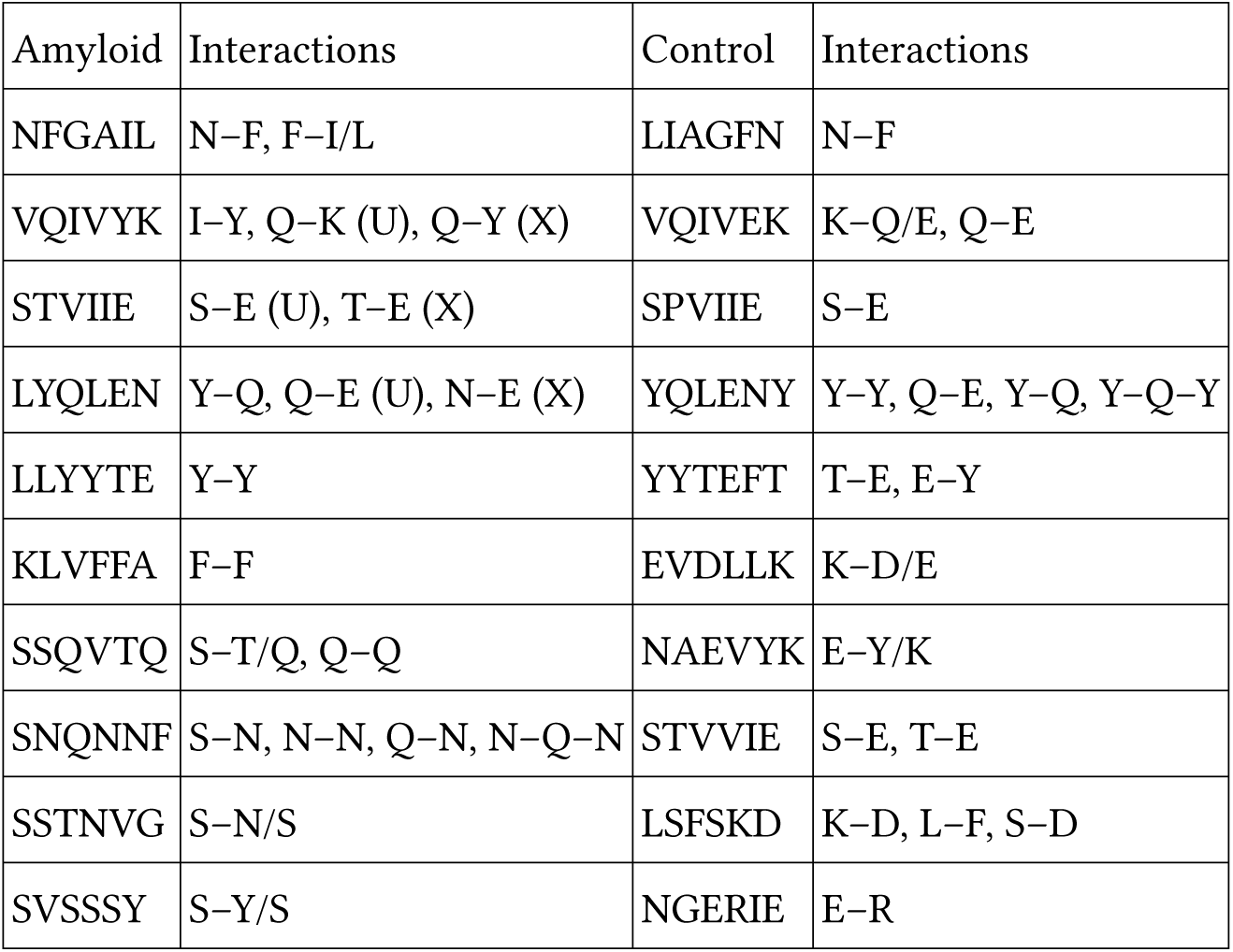
Intramolecular interactions between pair of amino acid residues in some of the low energy structures of hexapeptide monomers as visualised from the disconnectivity graphs given in Supplementary Information. The symbols X and U in parentheses represent the interactions present in helical/partial helical and hairpin structures, respectively. The aromatic rings (F/Y) may interact via a T-shaped or an offset stacked geometry. In SNQNNF, we also found three residues interacting simultaneously in the low energy structures for the monomer.

| Amyloid | Interactions | Control | Interactions |
| --- | --- | --- | --- |
| NFGAIL | N-F, F-I/L | LIAGFN | N-F |
| VQIVYK | I-Y, Q-K (U), Q-Y (X) | VQIVEK | K-Q/E, Q-E |
| STVIIE | S-E (U), T-E (X) | SPVIIE | S-E |
| LYQLEN | Y-Q, Q-E (U), N-E (X) | YQLENY | Y-Y, Q-E, Y-Q, Y-Q-Y |
| LLYYTE | Y-Y | YYTEFT | T-E, E-Y |
| KLVFFA | F-F | EVDLLK | K-D/E |
| SSQVTQ | S-T/Q, Q-Q | NAEVYK | E-Y/K |
| SNQNNF | S-N, N-N, Q-N, N-Q-N | STVVIE | S-E, T-E |
| SSTNVG | S-N/S | LSFSKD | K-D, L-F, S-D |
| SVSSSY | S-Y/S | NGERIE | E-R |

### Dimer heat capacity

Basin-hopping^45^ with rigid body moves and subsequent all-atom relaxation was used to sample the energy landscape for dimers. The low temperature heat capacity feature is taken as the first peak or inflection point above 0.086 kcal mol*^−^*^1^, as for the monomers (Fig. 1b). The structures within 2.6 kcal mol*^−^*^1^ of the global minimum are used in the end-to-end distance versus dihedral correlation plot. Interestingly, for amyloid hexapeptides, the structures with relatively large end-to-end distances and large positive dihedrals contribute to the C*_V_* feature of interest (Fig. 4b). For VQIVYK, very large negative dihedrals are also observed along with very large positive dihedrals. When the dihedral angles are close to 180 degrees (positive or negative), the corresponding structures are similar. However, for LYQLEN and YQLENY, the points in the correlation plot are close to each other. We note that LYQLEN forms amyloid at low pH,^85^ whereas the peptides are studied at neutral pH in the current study. NLGPVL is a control peptide that exhibits extended structures contributing to the C*_V_* peak of interest. However, further structural analysis reveals that the interstrand separation between monomers is larger compared to other amyloid forming dimers exhibiting extended conformations (Fig. 3). This result may be rationalised by taking into account the contribution from the L residue in the second position and P residue in NLGPVL. We suggest that these residues may not allow the strands to interact very closely and hence play a role in preventing aggregation of this peptide.

### Correlation between heat capacity and amyloid formation predictors

The correlation plot (Fig. 5) between the temperature at which the low temperature feature occurs in the C*_V_* plot and the propensity for amyloid formation as predicted using Aggrescan,^37^ shows that there is little overall correlation between the two quantities. However, further analysis reveals that the amyloid and control sequences derived from the same protein can probably be distinguished by comparing the temperature at which the first feature (excluding melting) occurs in the heat capacity plot between 0.076–0.300 kcal mol*^−^*^1^. The amyloid-forming sequences exhibit features at lower temperature compared to sequences with lower propensity for amyloid formation and the respective control sequences. This trend is evident when we compare the sequences within the sets derived from the same protein, such as tau (VQIVYK, VQIVEK, and NAEVYK), amylin (NFGAIL and NAGAIL), *β*_2_-microglobulin (LLYYTE, YYTEFT and EVDLLK), and A-*β* (KLVFFA and MVGGVV). The control sequences KAFIIQ, NLGPVL, and YQLENY lack such low temperature features. Another predictor that can be useful to estimate amyloid formation propensity is CamSol^86^. CamSol is designed to predict the intrinsic solubility of proteins. Aggregation-prone sequences exhibit lower solubility. Heptapeptides are the minimal sequences for which solubility can be obtained using CamSol. To obtain an approximate idea of solubility for the capped hexapeptides, we added alanine residues at the termini of the peptide sequences. As expected, the solubility exhibits a strong negative correlation with the aggregation propensity predicted using Aggrescan (Supplementary Information). Hence, the low temperature heat capacity feature is positively correlated with intrinsic solubility for peptide sequences occurring in the same context, i.e., within the same protein (Supplementary Information). As for Aggrescan there is little overall correlation between the two quantities, intrinsic solubility and low temperature C*_V_* feature. We note that the CamSol and Aggrescan predictors may have some intrinsic limitations for such small sequences. Both of them associate the control sequence KAILFL with high aggregation propensity, and the amyloid sequences with two or more serine residues are predicted to have high solubility and lower aggregation propensity. Overall, low temperature heat capacity features for proteins do not exhibit a strong correlation with aggregation propensity. However, they may be useful to compare the properties of short sequences occurring within the same protein, i.e, within the same context.

## **IV.** CONCLUSIONS

Heat capacity analysis of peptide monomers and dimers may provide insight into collective phenomena, such as aggregation and phase separation of proteins. We find that a variety of low energy structures with different backbone conformations contribute to the low temperature heat capacity feature for amyloid forming hexapeptide monomers. For control sequences, the C*_V_* peak of interest does not correspond to such a diverse set of structures. The structural competition can be diagnosed by a combination of geometric parameters, such as end-to-end distance and a dihedral angle. The heat capacity analysis for dimer conformations reveals that the amyloid forming hexapeptides preferentially contribute to the peak of interest via low energy conformations with large end-to-end distance and large positive dihedrals. The exact temperature at which the C*_V_* feature occurs is lower for the peptides with higher propensity for amyloid formation compared to control sequences derived from the same protein. However, this trend does not hold if we compare amyloidogenic hexapeptides from one protein and control sequences from a different protein.

We do not expect to extract a universal aggregating propensity from monomer and dimer properties. However, the analysis of low temperature heat capacity features reveals new opportunities for investigating and predicting collective behaviour. The distinctions we have identified for amyloid-forming hexapeptides, compared to controls with very similar sequences, suggest that further investigation could be fruitful. The fact that the current analysis could distinguish amyloid and control sequences with the same amino acid residues in reverse order, as for NFGAIL and LIAGFN, further illustrates how the energy landscape framework can be used to understand the context-dependent properties of amino acid residues in proteins.

## AUTHOR CONTRIBUTIONS

Nicy conceived the idea and designed the study with the help of D.J.W. Nicy performed the simulations and wrote the first draft. Both the authors analysed and interpreted the data, and edited the final version of the draft. D.J.W. supervised the project and developed all the original energy landscape software.

## FUNDING

This work was supported by Engineering and Physical Sciences Research Council (EPSRC) (D.J.W, grant number EP/N035003/1); the Cambridge Commonwealth, European and International Trust; the Allen, Meek and Read Fund; the Santander fund, St Edmund’s College, University of Cambridge; and the Trinity-Henry Barlow Honorary Award (Nicy).

## CONFLICT OF INTEREST

The authors declare no conflict of interest.

## DATA AVAILABILITY STATEMENT

The discrete path sampling databases can be obtained from authors upon request.

## Supporting information

Disconnectivity graphs

## Notes

### Competing Interest Statement

The authors have declared no competing interest.

