## Supplementary material for "Energy landscapes and heat capacity signatures for monomers and dimers of amyloid forming hexapeptides": Disconnectivity graphs

(Dated: 18 May 2023)

[a)]Electronic mail:

### SUPPLEMENTARY INFORMATION

| Hexapeptide | Peak/Inflection(*) temperatures | | | |
|---|---|---|---|---|
| **NFGAIL** | 0.076* | 0.364* | 0.701 | - |
| NAGAIL | 0.167* | 0.524 | - | - |
| LIAGFN | 0.043 | 0.223* | 0.392 | 0.962 |
| NLGPVL | 0.055 | 0.456 | - | - |
| **STVIIE** | 0.248 | 0.564 | - | - |
| STVIIP | 0.012 | 0.157 | 0.390 | 0.945 |
| SPVIIE | 0.030 | 0.295 | - | - |
| STVVIE | 0.101 | 0.235 | 0.947 | - |
| **VQIVYK** | 0.089 | 0.518 | 0.891 | - |
| VQIVEK | 0.011 | 0.131* | 0.269 | 1.031 |
| NAEVYK | 0.047 | 0.216 | 0.898 | - |
| **LYQLEN** | 0.079* | 0.227 | 0.385 | 0.802 |
| YQLENY | 0.024 | 0.281 | - | - |
| **LLYYTE** | 0.041 | 0.178 | 0.316 | - |
| YYTEFT | 0.017 | 0.242 | 0.427* | 1.145 |

TABLE S1: Hexapeptides, and $k_B T$ ($\mathrm{kcal\,mol^{-1}}$) at which peaks or distinct inflection points are observed. The inflection points are marked with an asterisk (*).

| Hexapeptide | Peak/Inflection(*) temperatures | | | |
|---|---|---|---|---|
| GAIIGL | 0.219* | 0.598 | - | - |
| GGVVIA | 0.034 | 0.377* | 0.582 | - |
| MVGGVV | 0.037 | 0.213 | 0.630 | - |
| GAILSS | 0.179 | 0.310 | - | - |
| SSQVTQ | 0.218 | 0.426 | - | - |
| SSTNVG | 0.054 | 0.241 | - | - |
| SNQNNF | 0.016 | 0.199 | 0.312 | 1.011 |
| SVSSSY | 0.039 | 0.240 | 0.514 | - |
| GYVIIK | 0.016 | 0.437 | - | - |
| KLVFFA | 0.150 | 0.439 | - | - |
| KAFIIQ | 0.259 | - | - | - |
| KAILFL | 0.112 | 0.309 | - | - |
| EVDLLK | 0.021 | 0.204* | 0.417 | - |
| LSFSKD | 0.011 | 0.101* | 0.368 | 0.860 |
| NGERIE | 0.155 | 0.404 | - | - |

TABLE S2: Hexapeptides, and $k_\mathrm{B}T$ ( $\mathrm{kcal\,mol^{-1}}$) at which peaks or distinct inflection points are observed. The inflection points are marked with an asterisk (*).

| Hexapeptide | Peak/Inflection(*) temperatures | | | |
|---|---|---|---|---|
| **NFGAIL** | 0.060 | $0.432^{*}$ | 0.859 | - |
| NAGAIL | 0.016 | 0.107 | 1.148 | - |
| LIAGFN | 0.167 | $0.294^{*}$ | 1.080 | - |
| NLGPVL | 0.100 | $0.458^{i}$ | 1.095 | - |
| **STVIIE** | 0.122 | 0.540 | $1.096^{i}$ | - |
| STVIIP | 0.036 | 0.547 | 1.051 | - |
| SPVIIE | 0.023 | 0.229 | 0.400 | - |
| STVVIE | 0.043 | 0.354 | 0.667 | - |
| **VQIVYK** | $0.081^{*}$ | 0.268 | 1.123 | - |
| VQIVEK | 0.061 | 0.420 | 0.891 | - |
| NAEVYK | 0.107 | 0.718 | - | - |
| **LYQLEN** | 0.042 | $0.173^{*}$ | 0.600 | 0.888 |
| YQLENY | $0.271^{*}$ | 0.408 | - | - |
| **LLYYTE** | $0.021^{i}$ | 0.281 | 0.902 | - |
| YYTEFT | 0.057 | 0.215 | 0.418 | $1.156^{i}$ |

TABLE S3: Dimers of hexapeptides, and $k_{\mathrm{B}}T$ ( $\mathrm{kcal\,mol^{-1}}$) at which peaks or distinct inflection points are observed. The inflection points are marked with an asterisk (*).

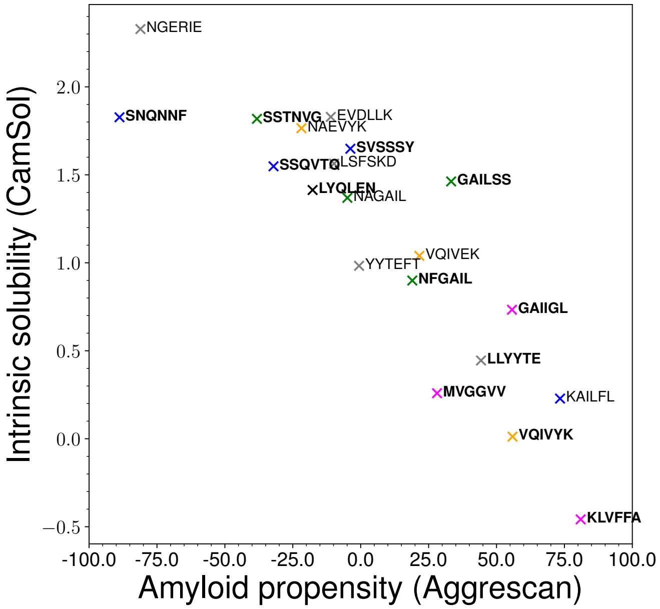


(a) Correlation plot between amyloid predictor Aggrescan and intrinsic solubility predictor CamSol.

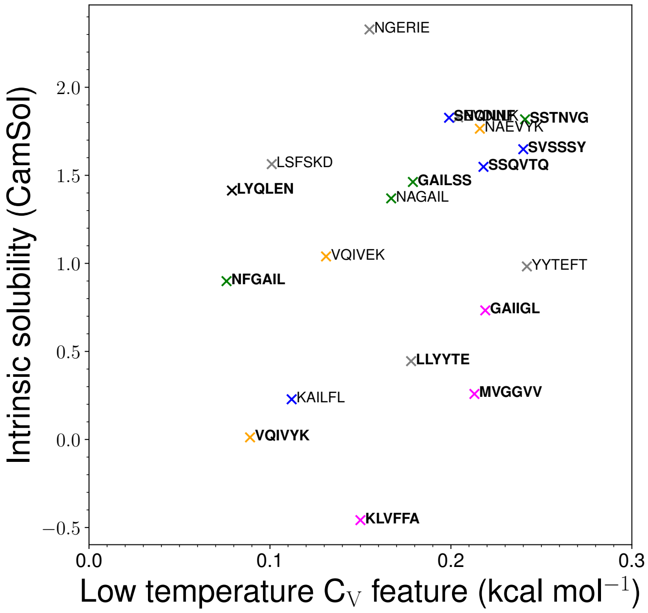


(b) Correlation plot between low temperature heat capacity feature and intrinsic solubility as predicted using CamSol.

FIG. S1: Correlation plots between different amyloid predictors.

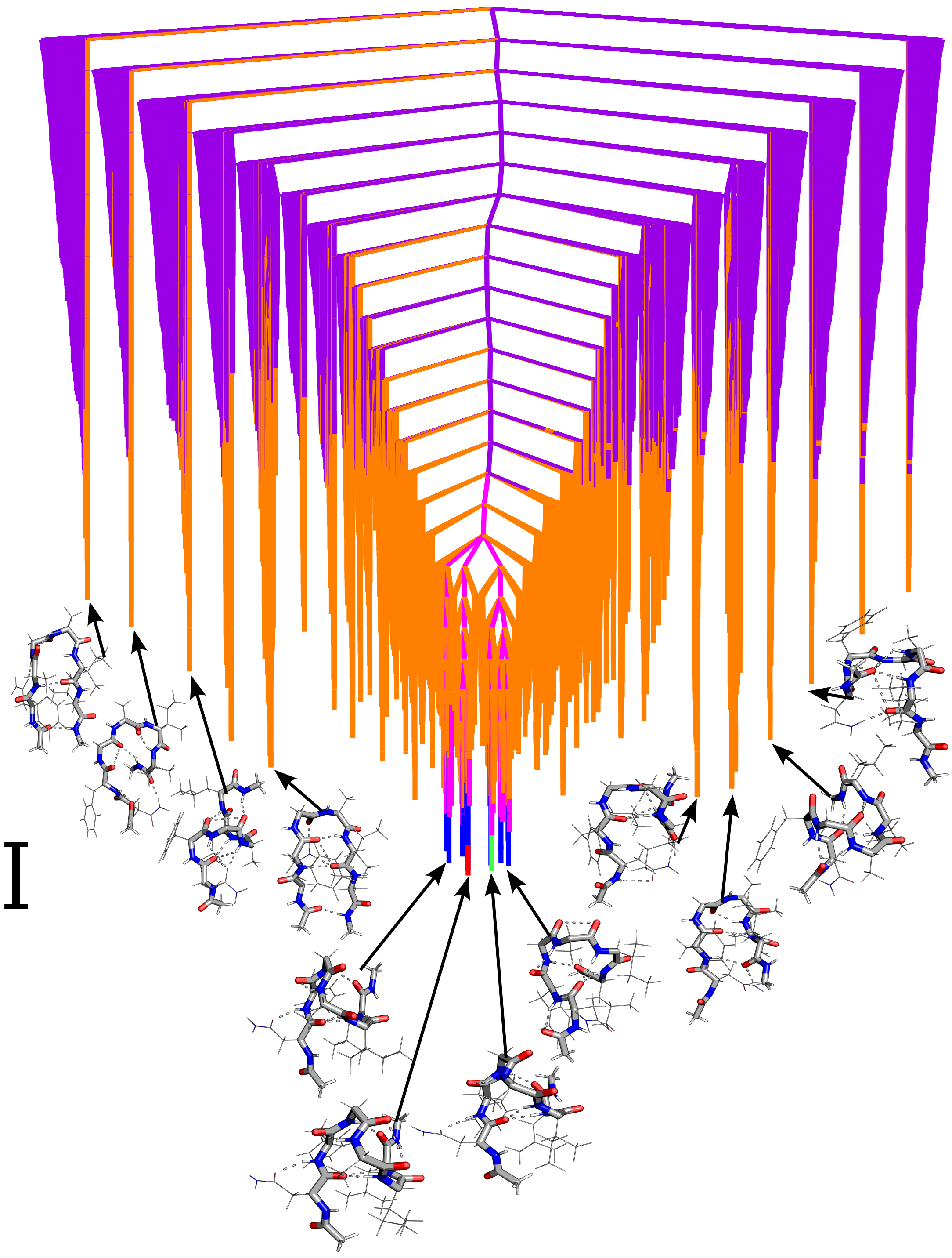

FIG. S2: Disconnectivity graphs for **NFGAIL** monomer. Local minima representing transition for peaks/inflection points are represented by red to blue (peak 1), green to orange (peak 2), pink to purple (peak 3) and grey to yellow (peak 4). The scalebar represents 1 kcal mol$^{-1}$.

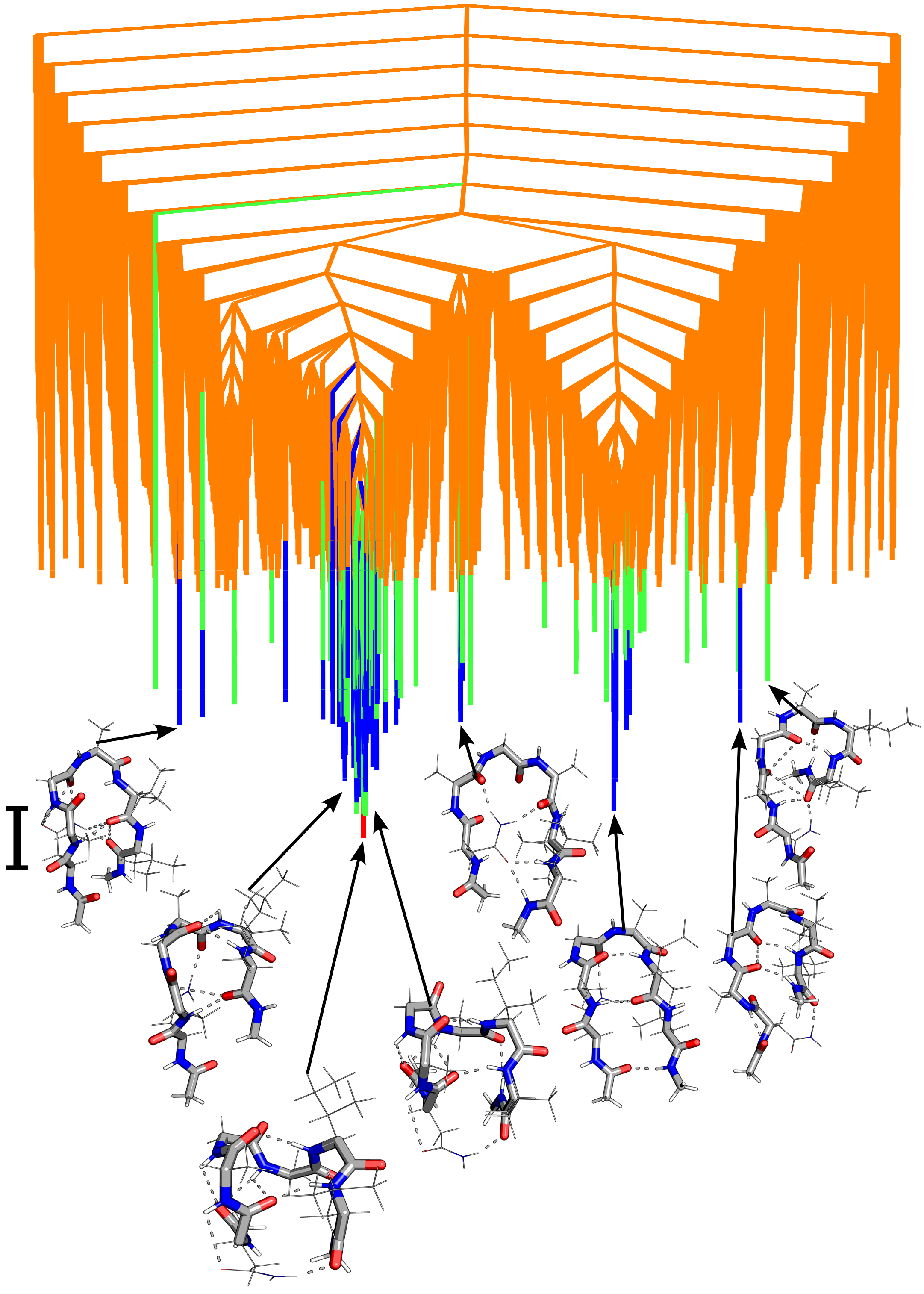

FIG. S3: Disconnectivity graphs for NAGAIL monomer. Local minima representing transition for peaks/inflection points are represented by red to blue (peak 1), green to orange (peak 2), pink to purple (peak 3) and grey to yellow (peak 4). The scalebar represents 1 kcal mol$^{-1}$.

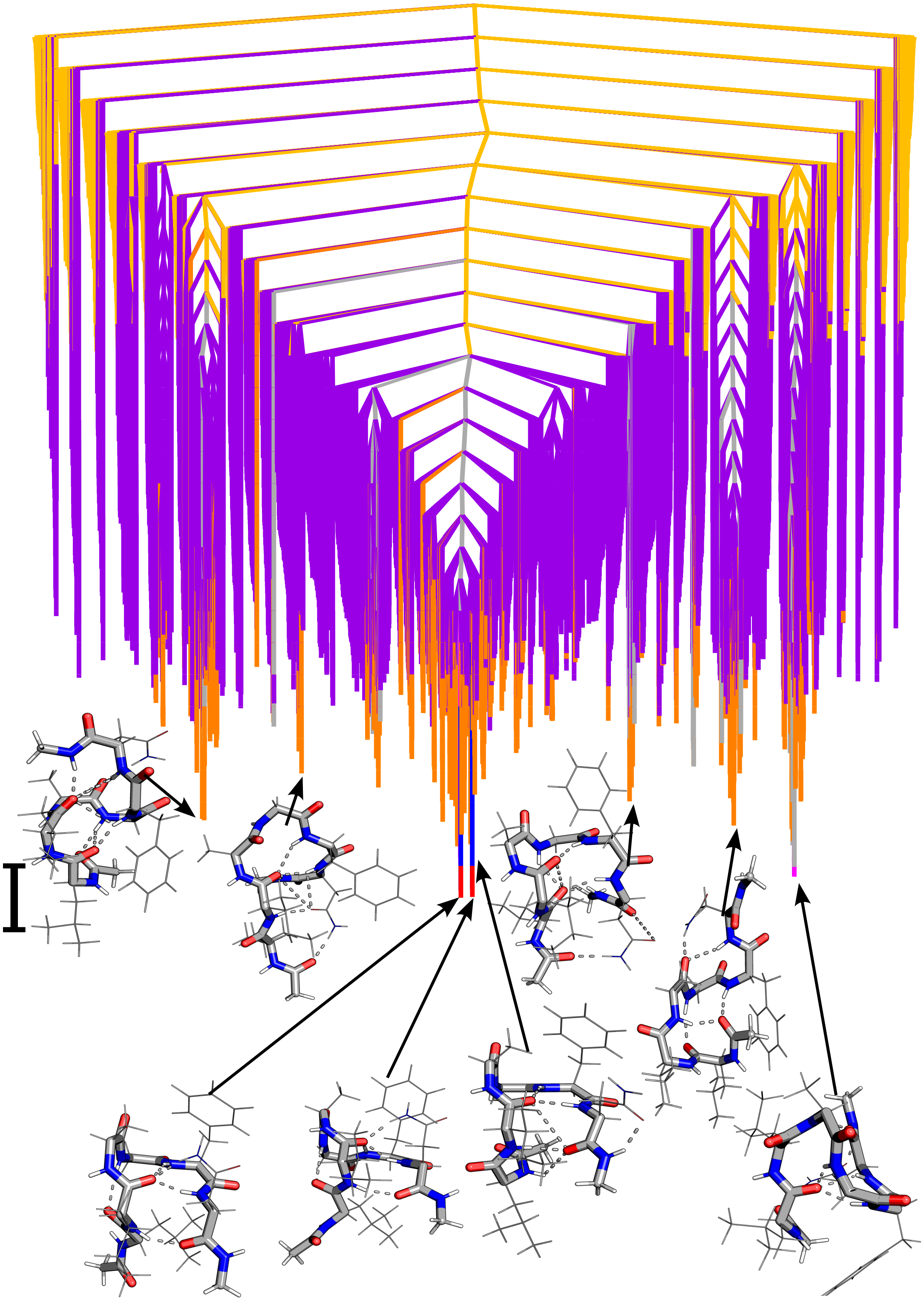

FIG. S4: Disconnectivity graphs for LIAGFN monomer. Local minima representing transition for peaks/inflection points are represented by red to blue (peak 1), green to orange (peak 2), pink to purple (peak 3) and grey to yellow (peak 4). The scalebar represents 1 kcal $mol^{-1}$.

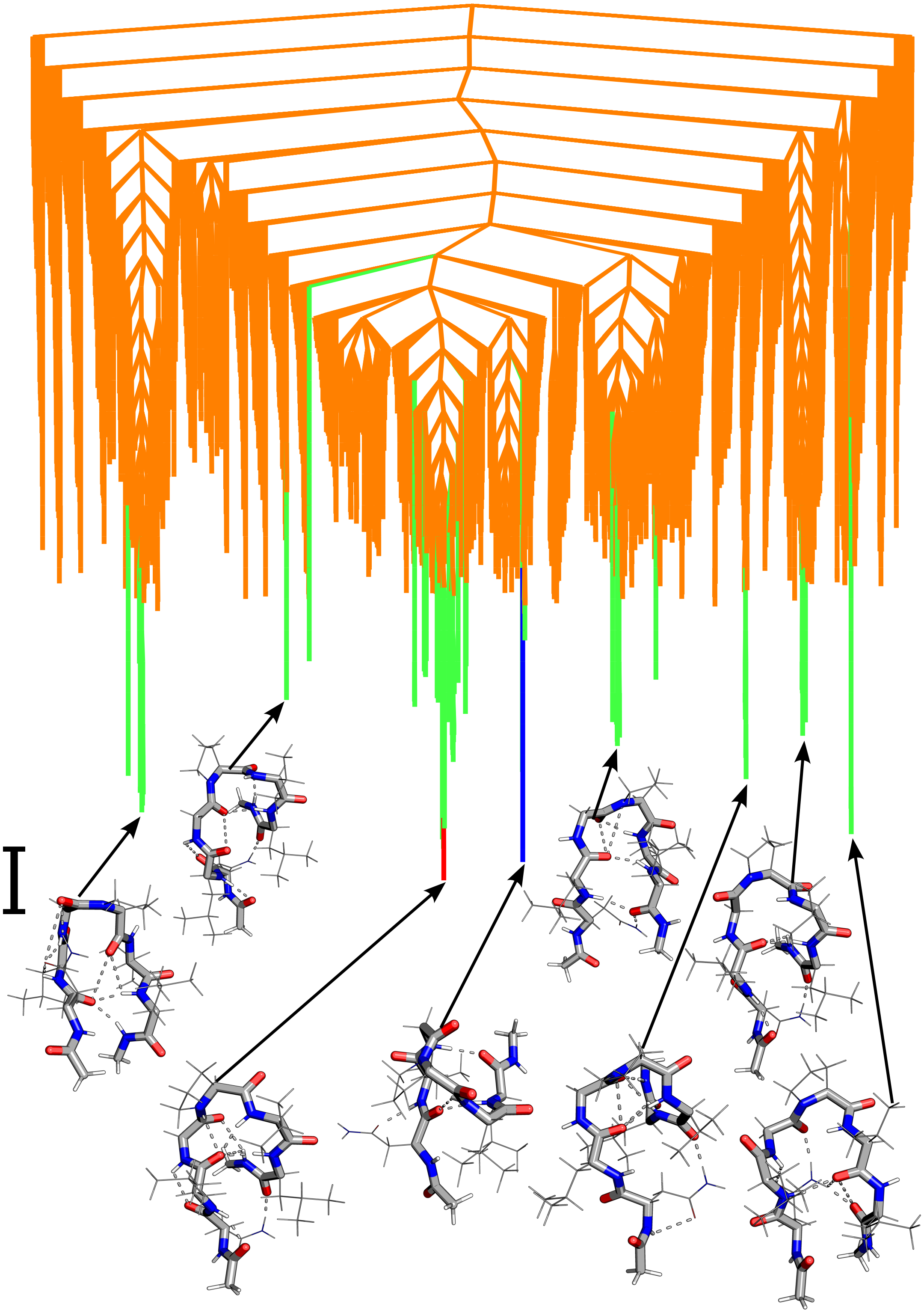

FIG. S5: Disconnectivity graphs for NLGPVL monomer. Local minima representing transition for peaks/inflection points are represented by red to blue (peak 1), green to orange (peak 2), pink to purple (peak 3) and grey to yellow (peak 4). The scalebar represents 1 kcal $mol^{-1}$.

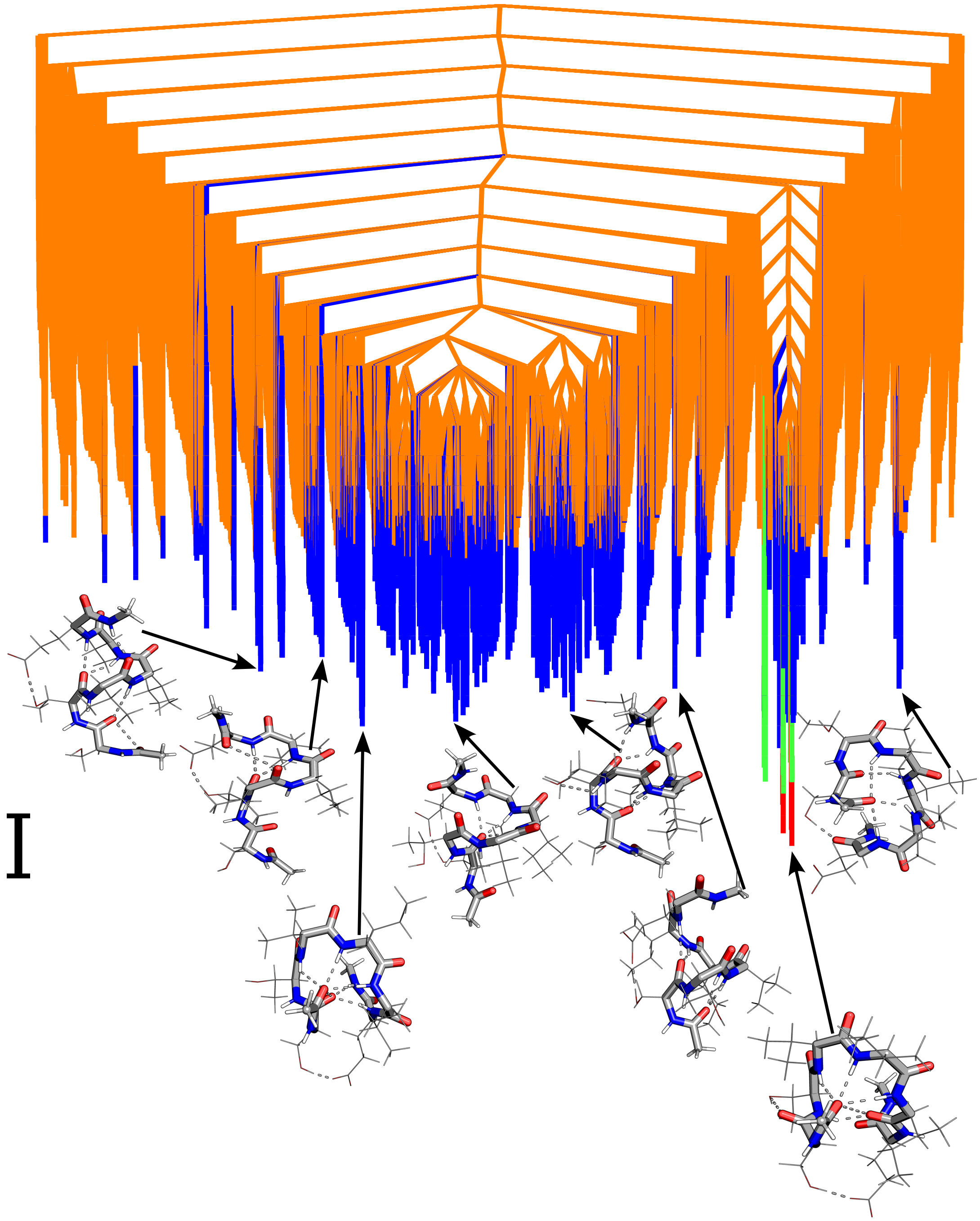

FIG. S6: Disconnectivity graphs for **STVIIE** monomer. Local minima representing transition for peaks/inflection points are represented by red to blue (peak 1), green to orange (peak 2), pink to purple (peak 3) and grey to yellow (peak 4). The scalebar represents 1 kcal mol$^{-1}$.

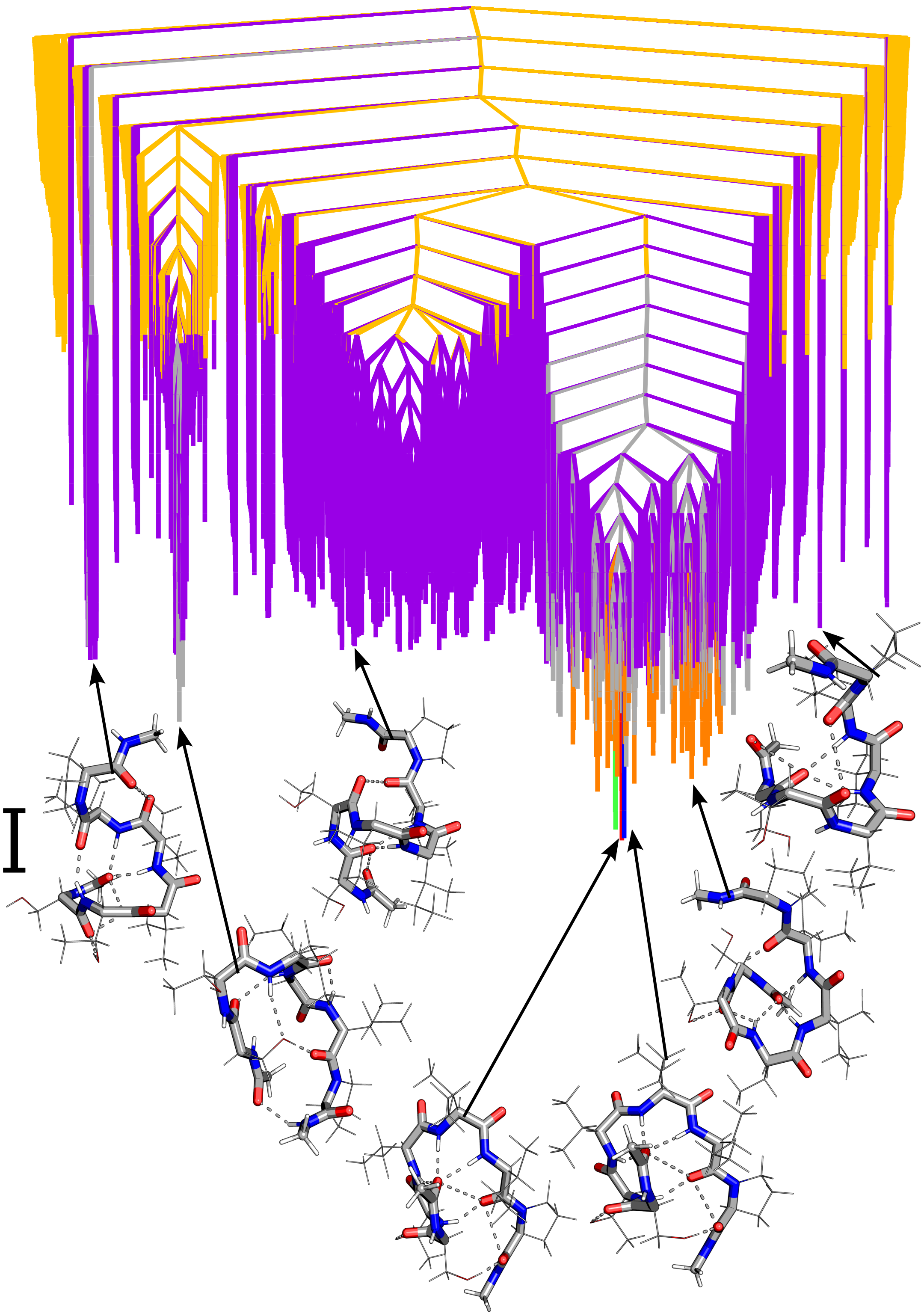

FIG. S7: Disconnectivity graphs for STVIIP monomer. Local minima representing transition for peaks/inflection points are represented by red to blue (peak 1), green to orange (peak 2), pink to purple (peak 3) and grey to yellow (peak 4). The scalebar represents 1 kcal mol$^{-1}$.

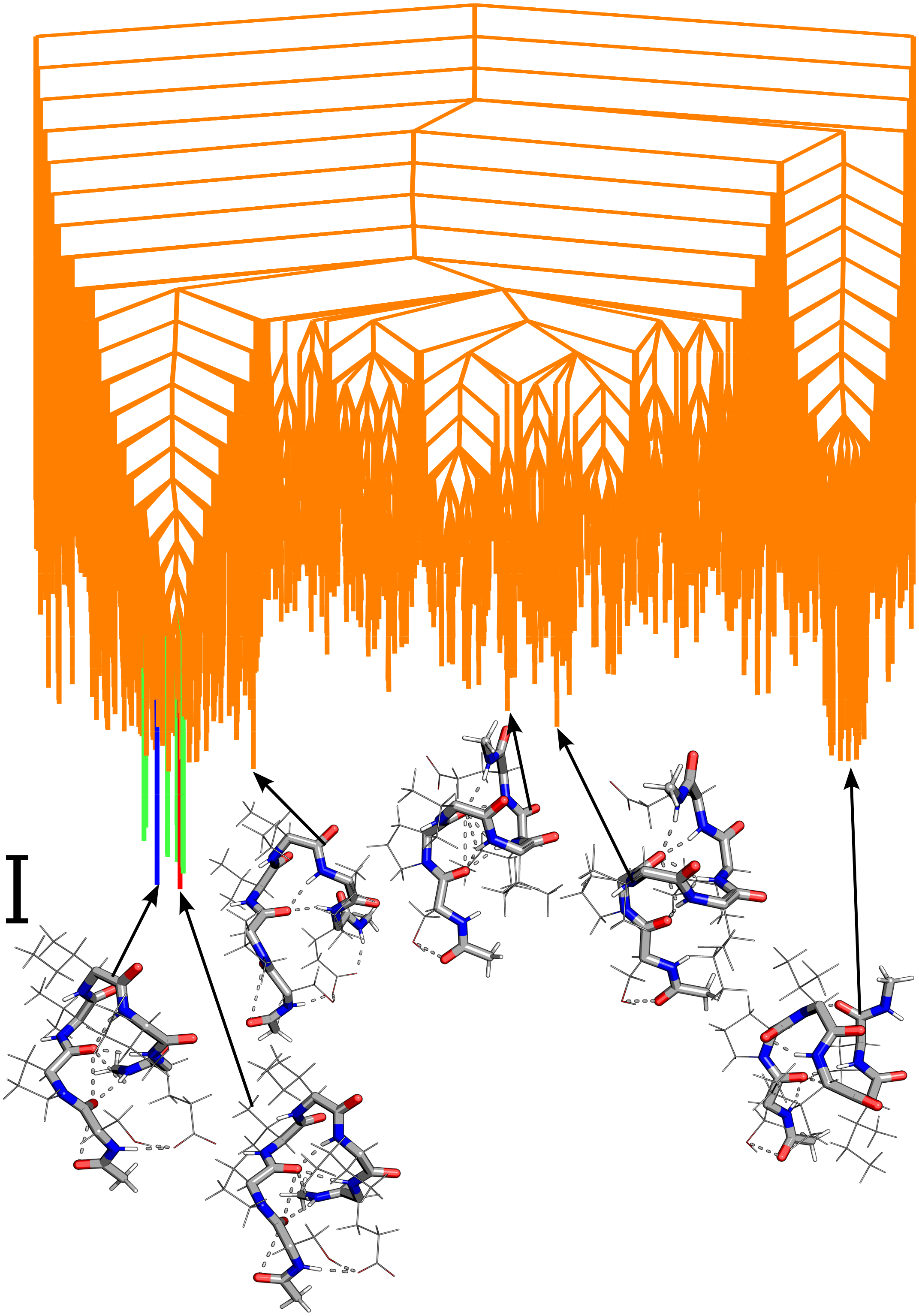

FIG. S8: Disconnectivity graphs for SPVIIE monomer. Local minima representing transition for peaks/inflection points are represented by red to blue (peak 1), green to orange (peak 2), pink to purple (peak 3) and grey to yellow (peak 4). The scalebar represents 1 kcal $mol^{-1}$.

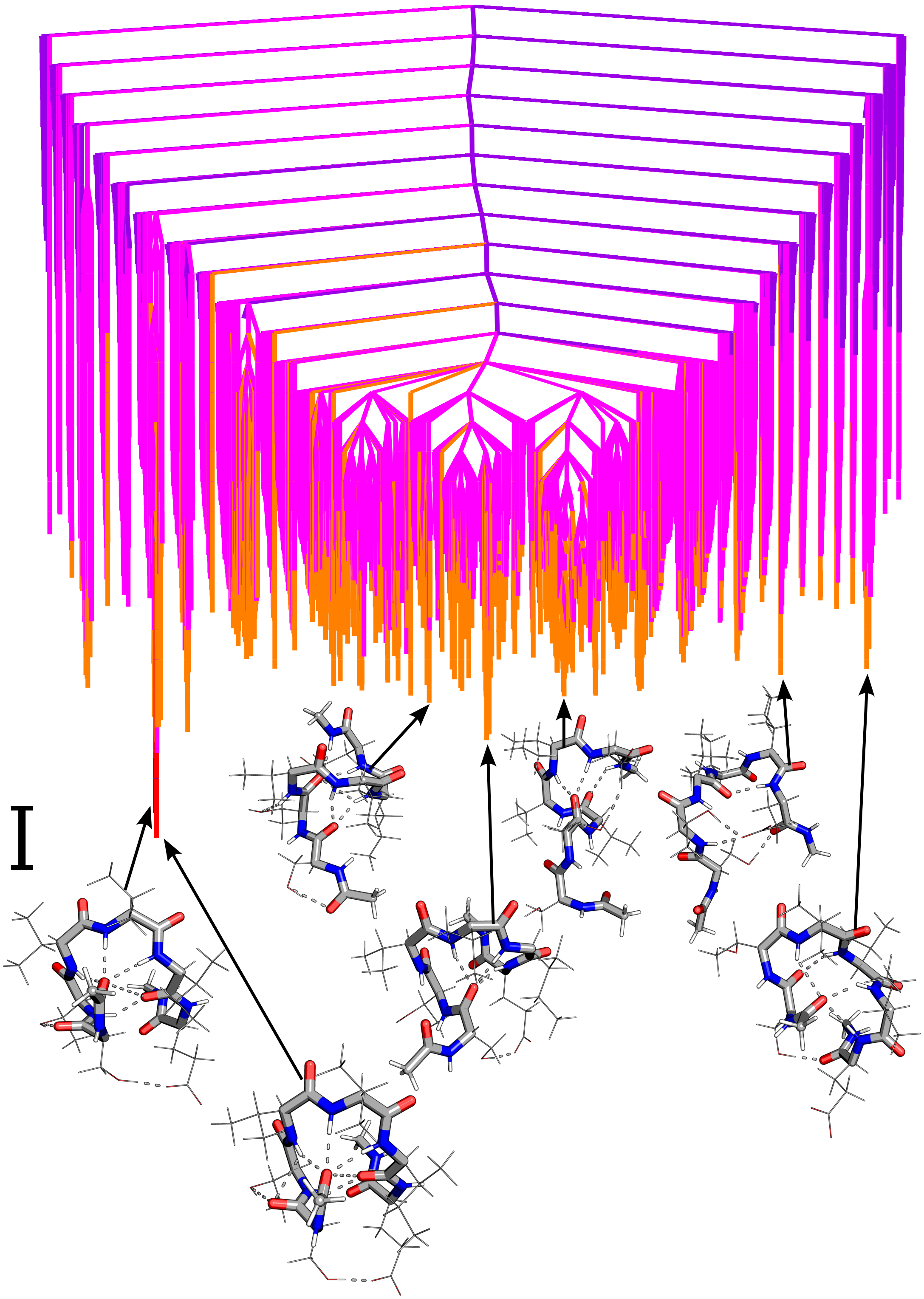

FIG. S9: Disconnectivity graphs for STVVIE monomer. Local minima representing transition for peaks/inflection points are represented by red to blue (peak 1), green to orange (peak 2), pink to purple (peak 3) and grey to yellow (peak 4). The scalebar represents 1 kcal mol$^{-1}$.

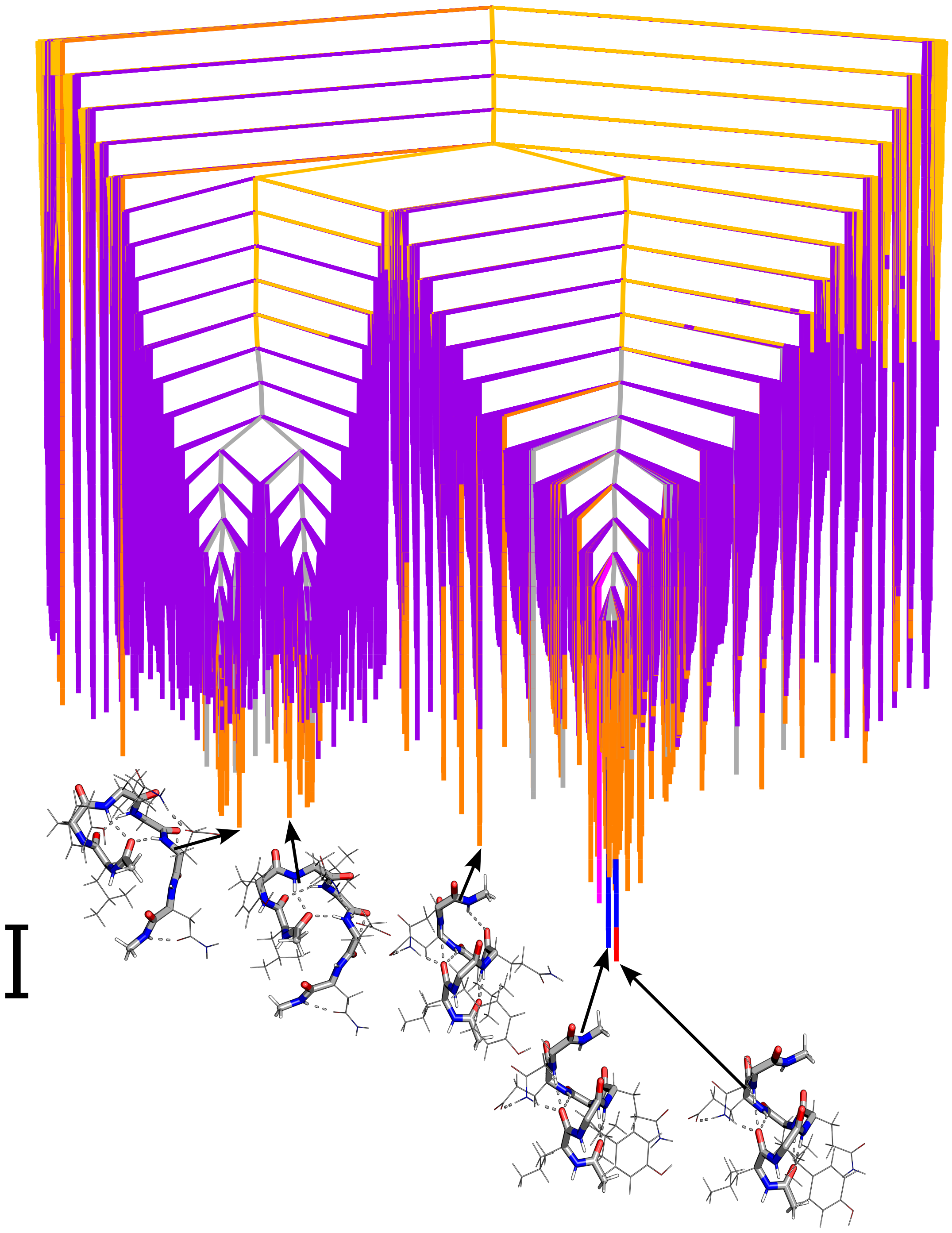

FIG. S10: Disconnectivity graphs for **LYQLEN** monomer. Local minima representing transition for peaks/inflection points are represented by red to blue (peak 1), green to orange (peak 2), pink to purple (peak 3) and grey to yellow (peak 4). The scalebar represents 1 kcal $mol^{-1}$.

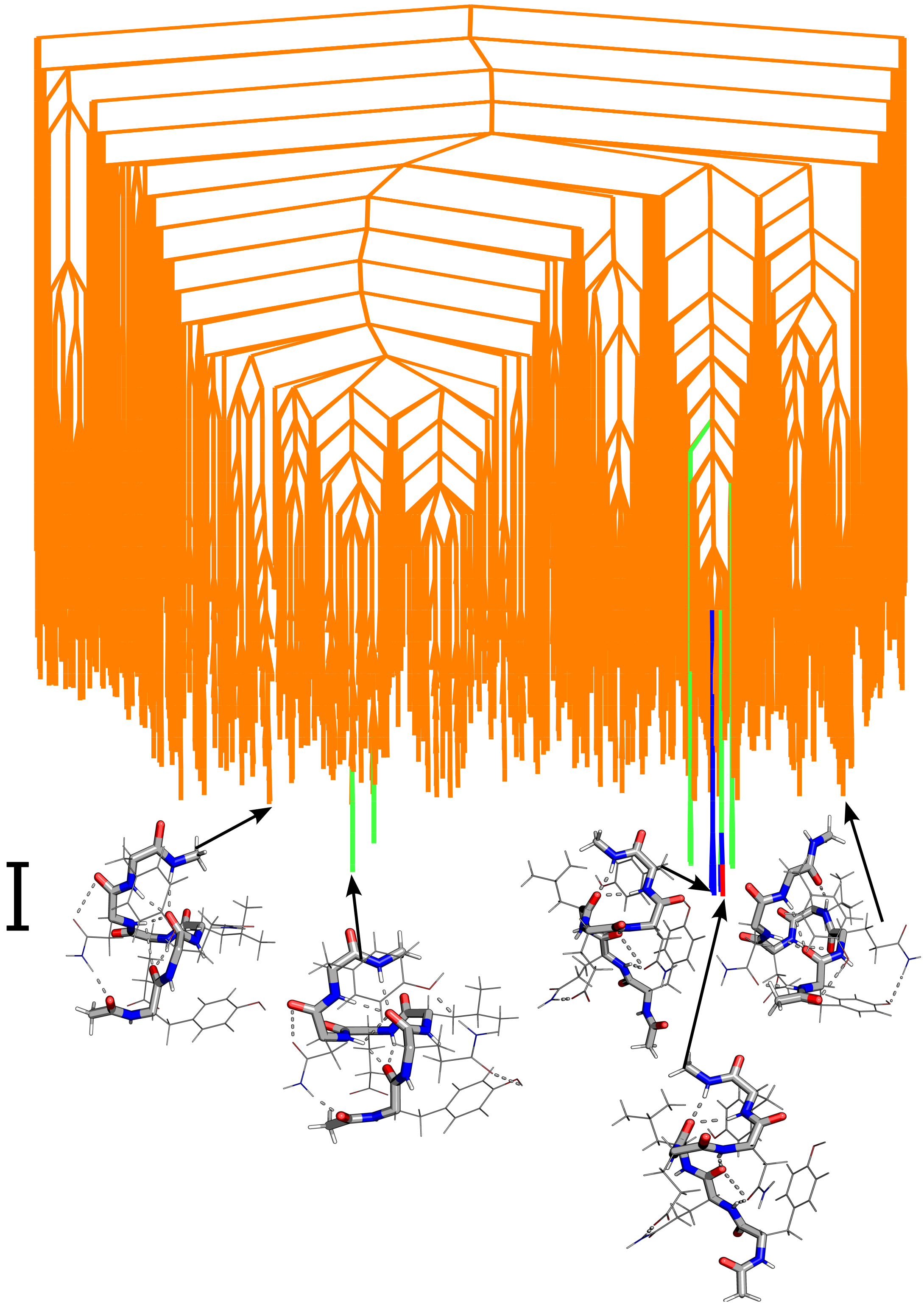

FIG. S11: Disconnectivity graphs for YQLENY monomer. Local minima representing transition for peaks/inflection points are represented by red to blue (peak 1), green to orange (peak 2), pink to purple (peak 3) and grey to yellow (peak 4). The scalebar represents 1 kcal mol$^{-1}$.

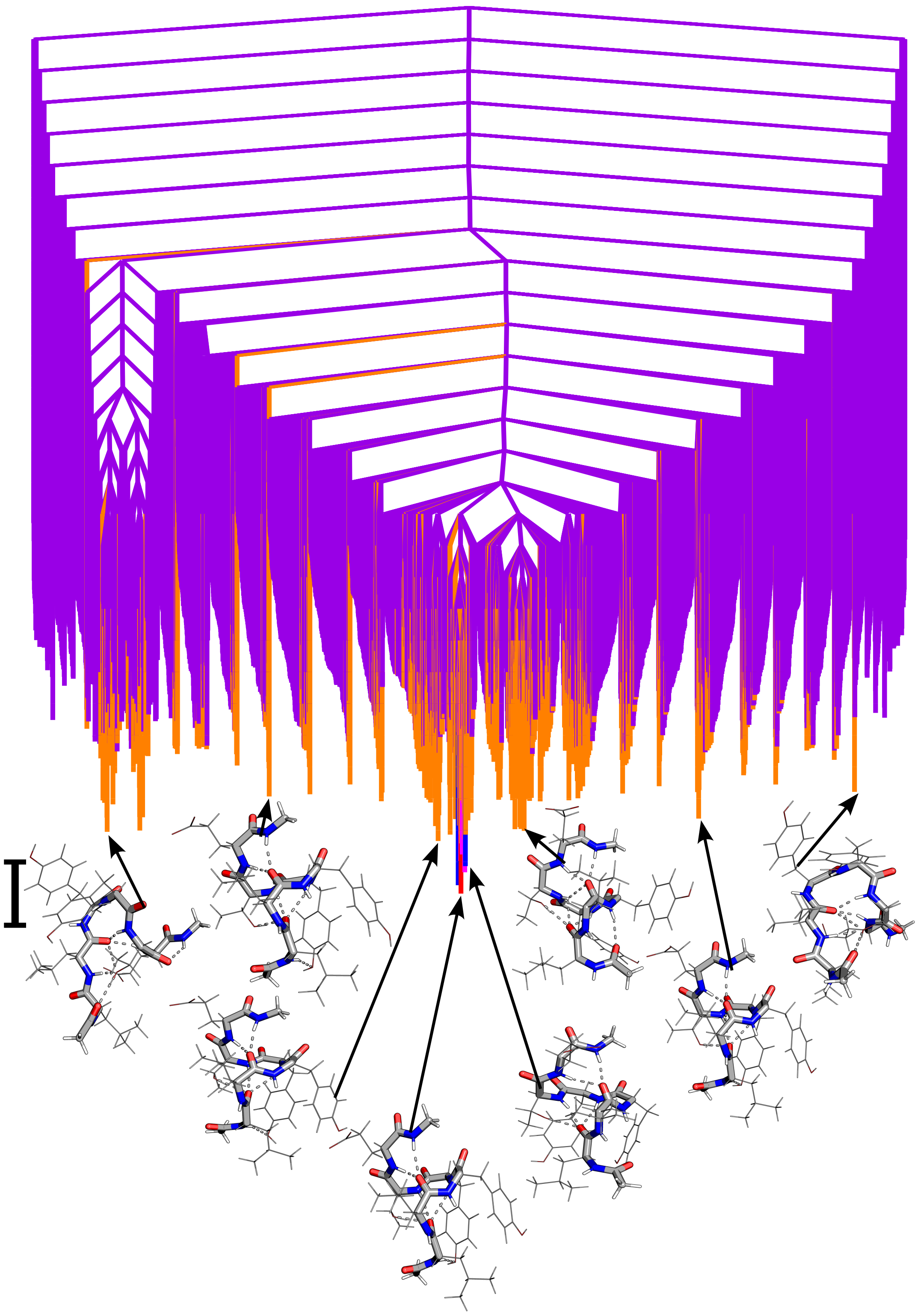

FIG. S12: Disconnectivity graphs for **LLYYTE** monomer. Local minima representing transition for peaks/inflection points are represented by red to blue (peak 1), green to orange (peak 2), pink to purple (peak 3) and grey to yellow (peak 4). The scalebar represents 1 kcal mol$^{-1}$.

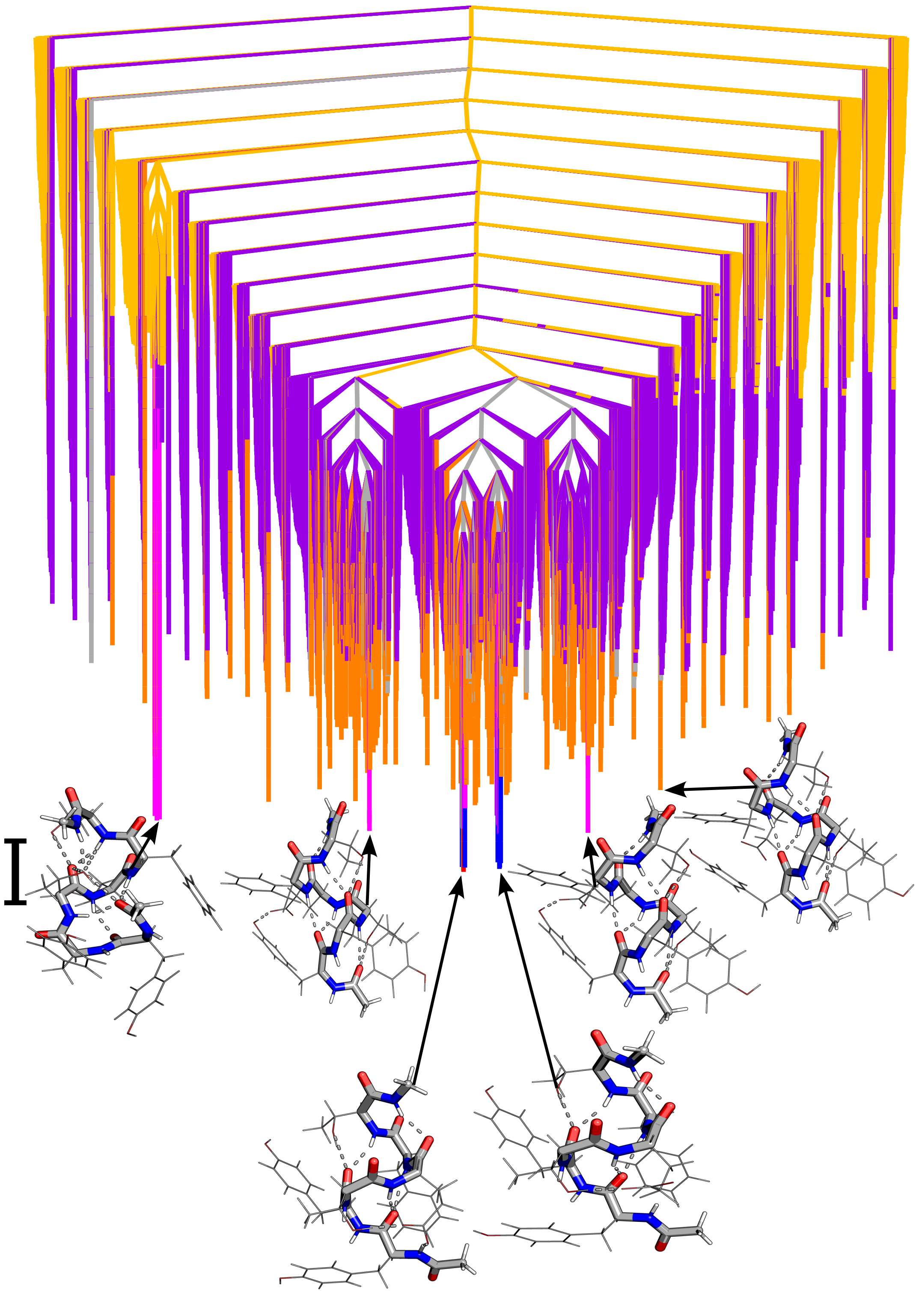

FIG. S13: Disconnectivity graphs for YYTEFT monomer. Local minima representing transition for peaks/inflection points are represented by red to blue (peak 1), green to orange (peak 2), pink to purple (peak 3) and grey to yellow (peak 4). The scalebar represents 1 kcal mol$^{-1}$.

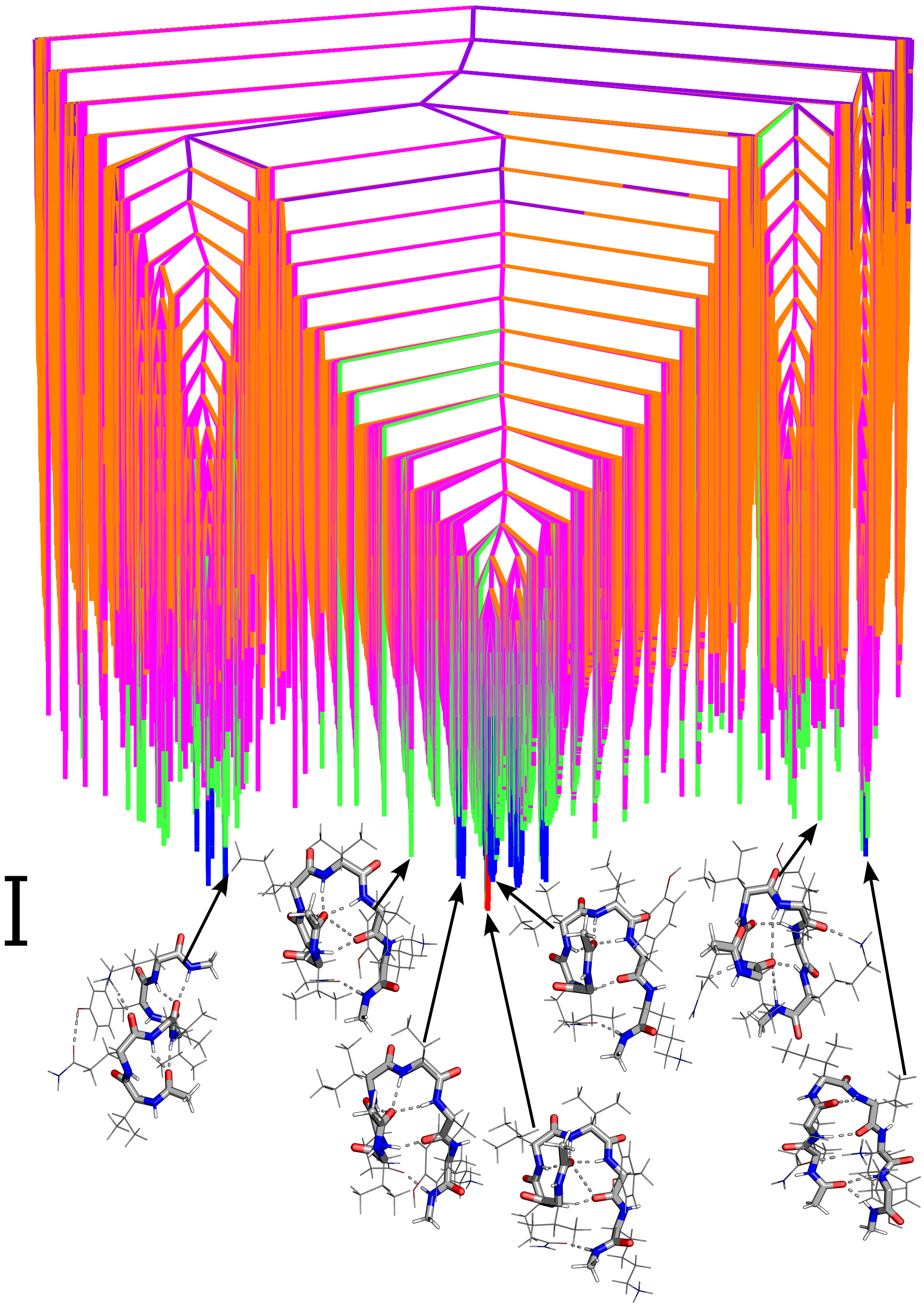

FIG. S14: Disconnectivity graphs for **VQIVYK** monomer. Local minima representing transition for peaks/inflection points are represented by red to blue (peak 1), green to orange (peak 2), pink to purple (peak 3) and grey to yellow (peak 4). The scalebar represents 1 kcal mol$^{-1}$.

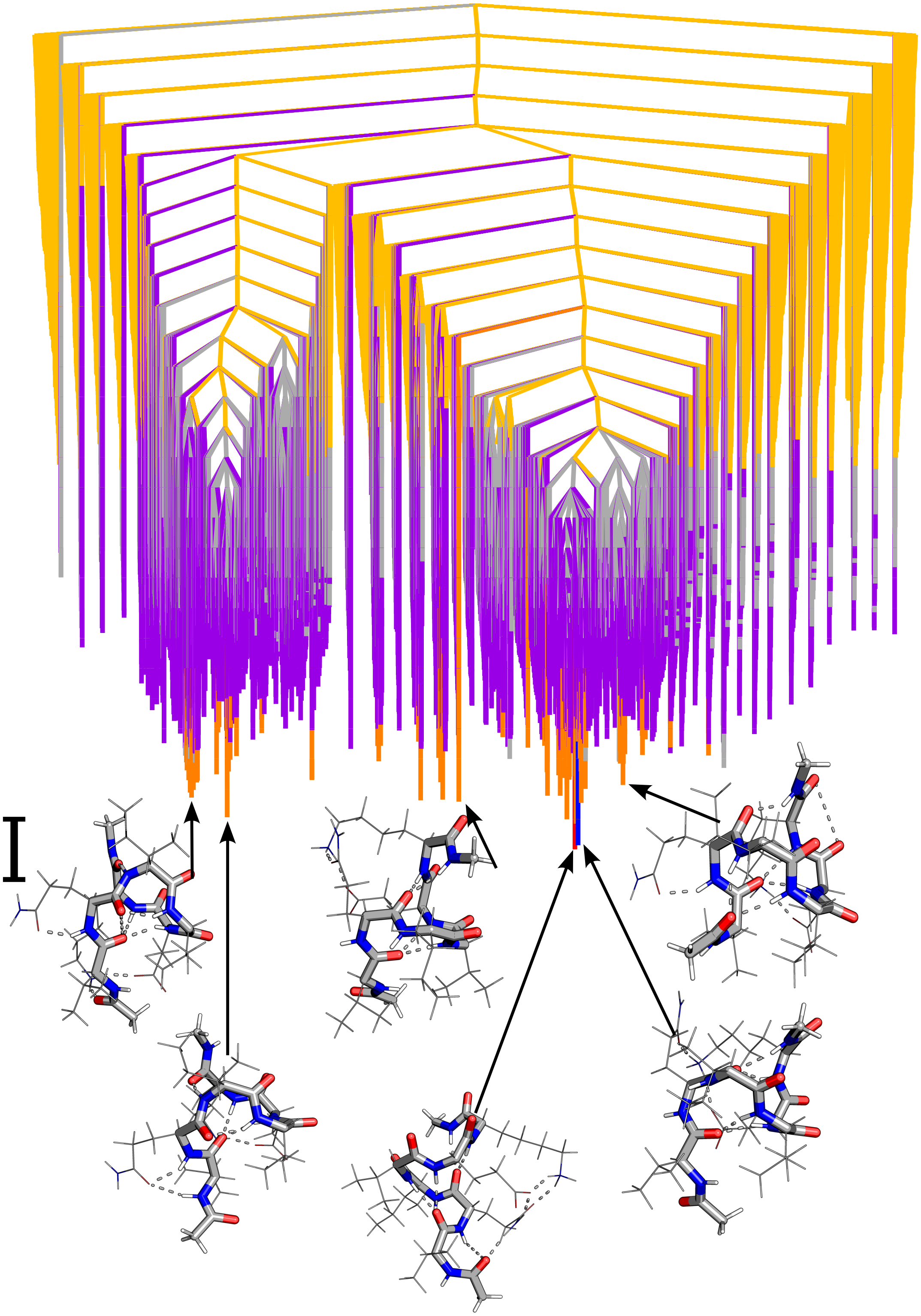

FIG. S15: Disconnectivity graphs for VQIVEK monomer. Local minima representing transition for peaks/inflection points are represented by red to blue (peak 1), green to orange (peak 2), pink to purple (peak 3) and grey to yellow (peak 4). The scalebar represents 1 kcal mol$^{-1}$.

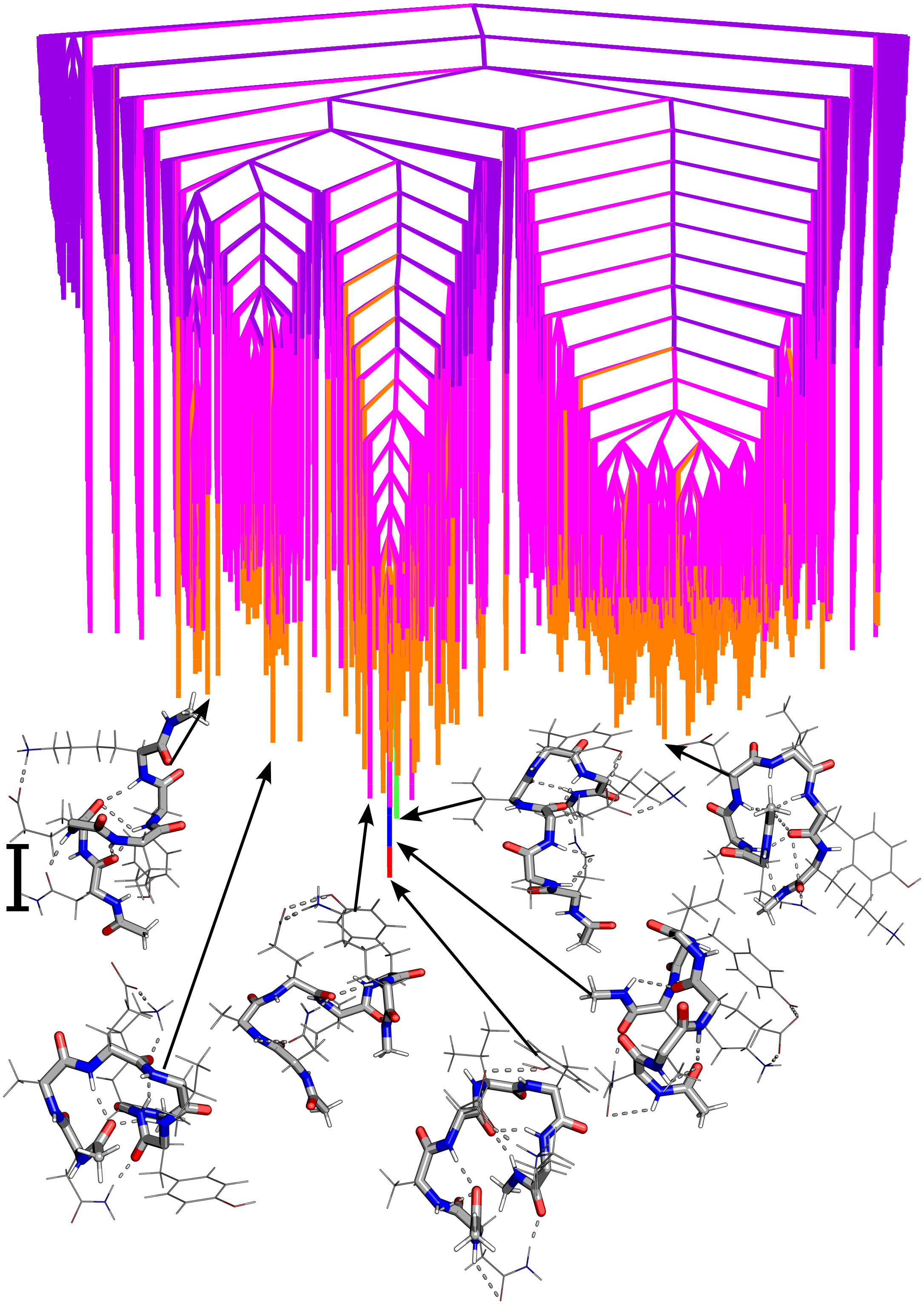

FIG. S16: Disconnectivity graphs for NAEVYK monomer. Local minima representing transition for peaks/inflection points are represented by red to blue (peak 1), green to orange (peak 2), pink to purple (peak 3) and grey to yellow (peak 4). The scalebar represents 1 kcal $mol^{-1}$.

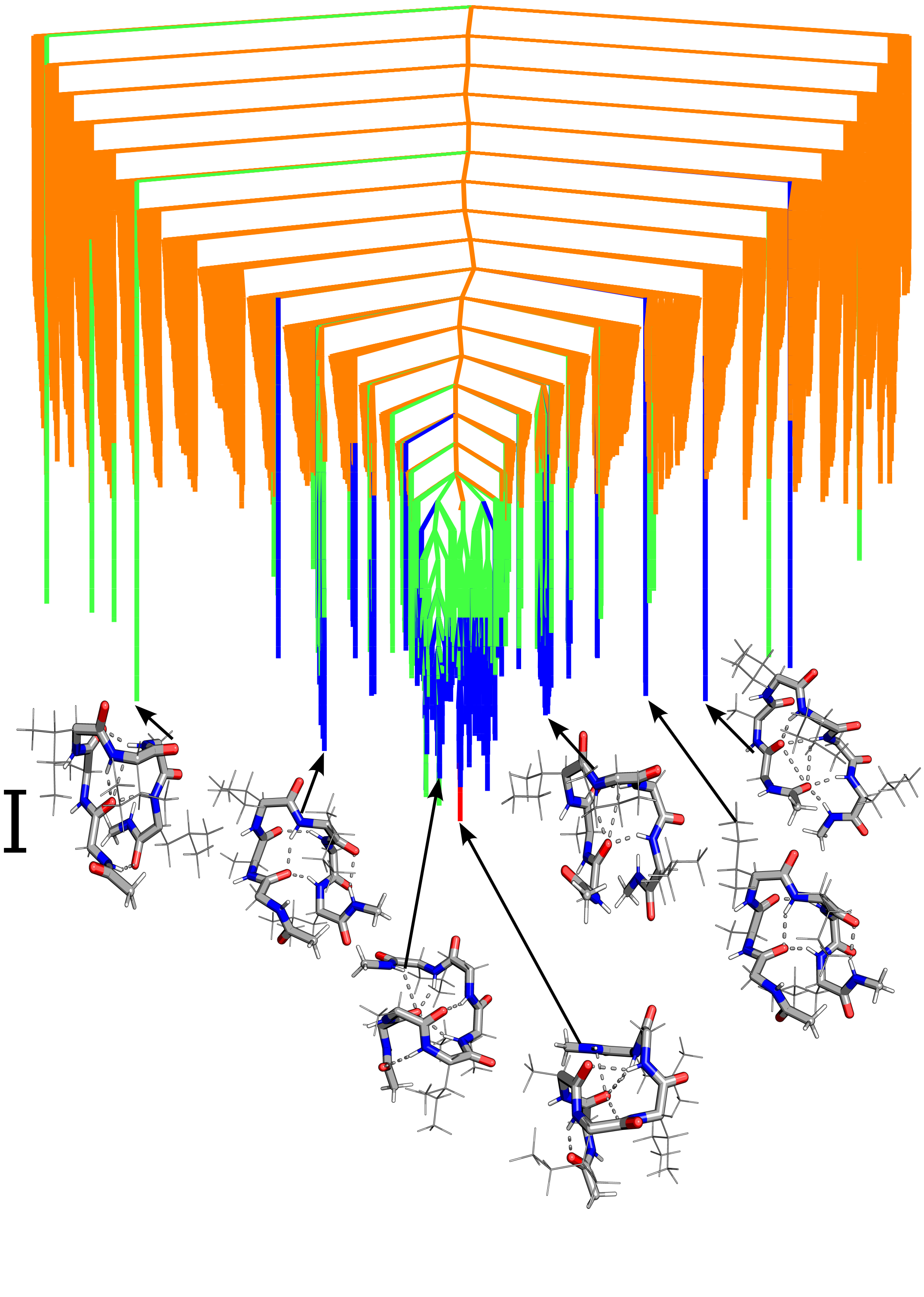

FIG. S17: Disconnectivity graphs for **GAIIGL** monomer. Local minima representing transition for peaks/inflection points are represented by red to blue (peak 1), green to orange (peak 2), pink to purple (peak 3) and grey to yellow (peak 4). The scalebar represents 1 kcal $mol^{-1}$.

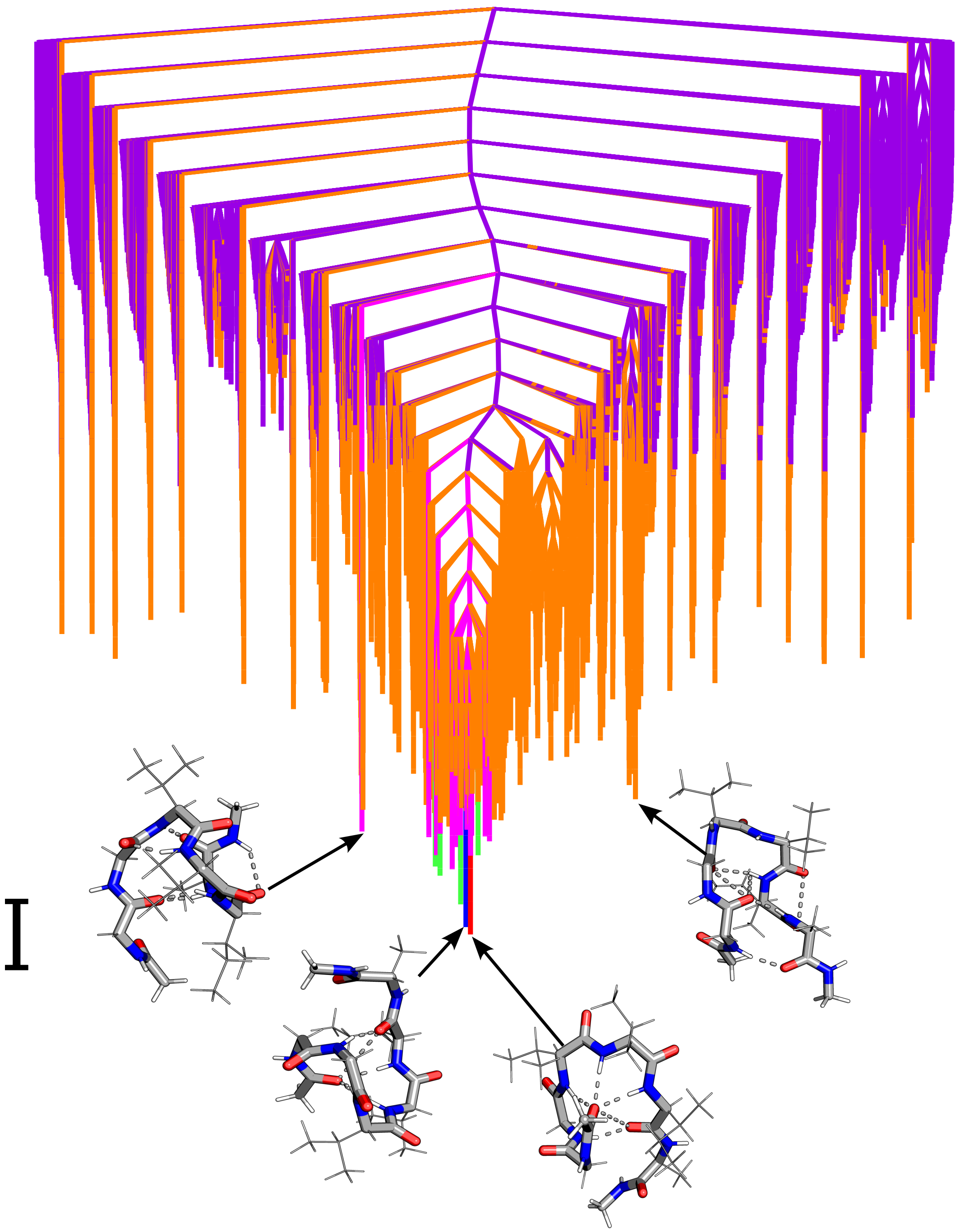

FIG. S18: Disconnectivity graphs for **GGVVIA** monomer. Local minima representing transition for peaks/inflection points are represented by red to blue (peak 1), green to orange (peak 2), pink to purple (peak 3) and grey to yellow (peak 4). The scalebar represents 1 kcal mol$^{-1}$.

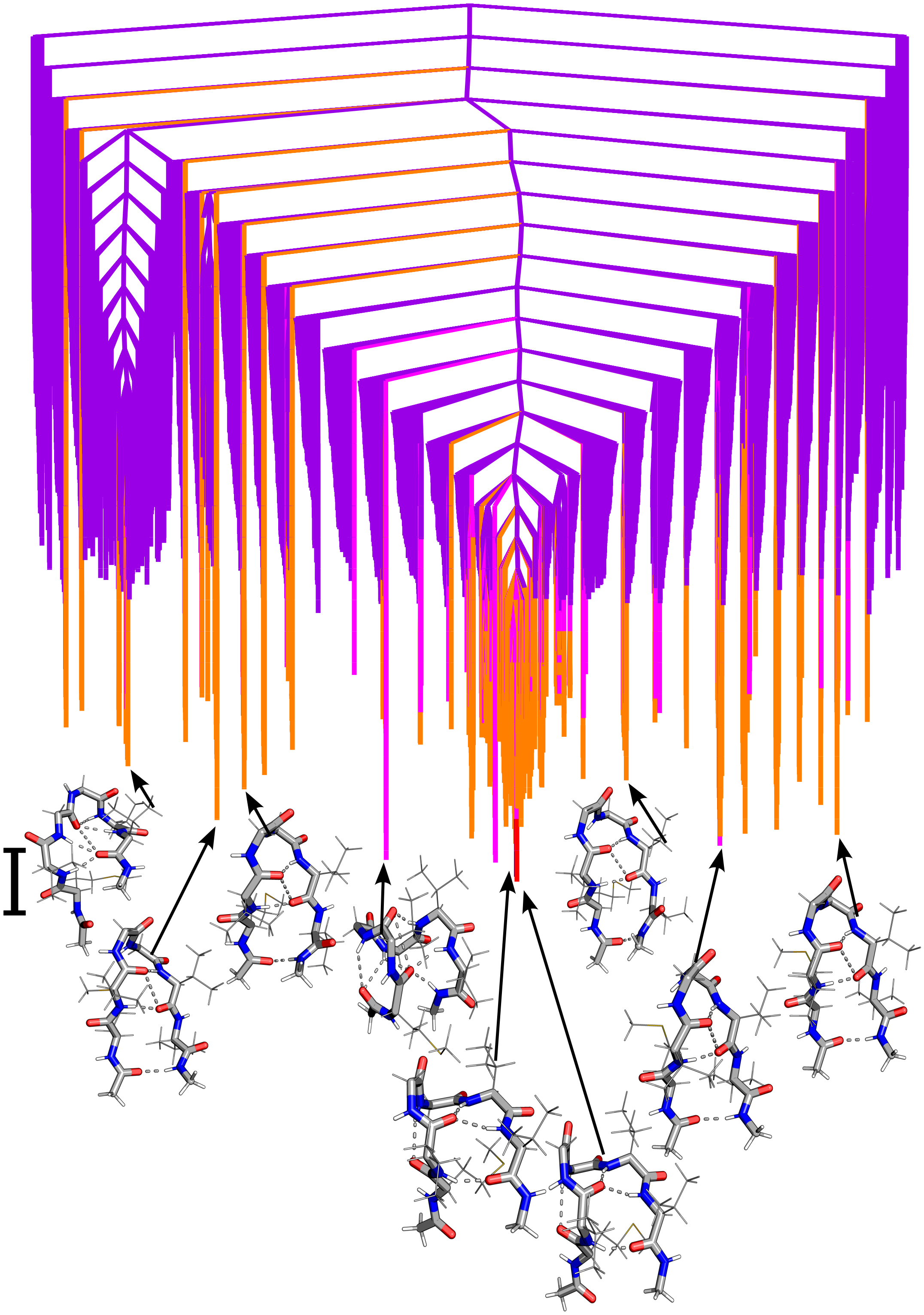

FIG. S19: Disconnectivity graphs for **MVGGVV** monomer. Local minima representing transition for peaks/inflection points are represented by red to blue (peak 1), green to orange (peak 2), pink to purple (peak 3) and grey to yellow (peak 4). The scalebar represents 1 kcal $mol^{-1}$.

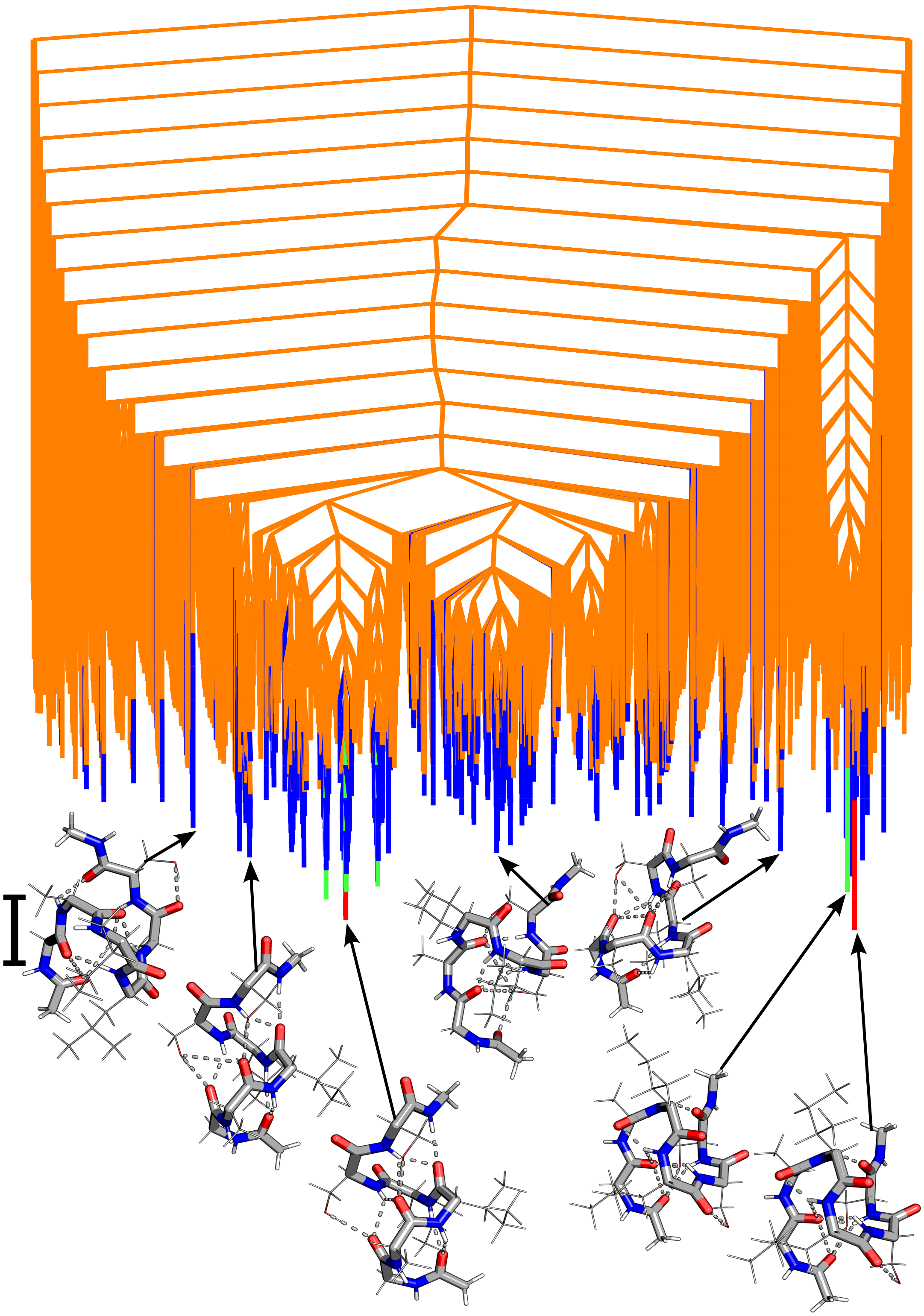

FIG. S20: Disconnectivity graphs for **GAILSS** monomer. Local minima representing transition for peaks/inflection points are represented by red to blue (peak 1), green to orange (peak 2), pink to purple (peak 3) and grey to yellow (peak 4). The scalebar represents 1 kcal mol$^{-1}$.

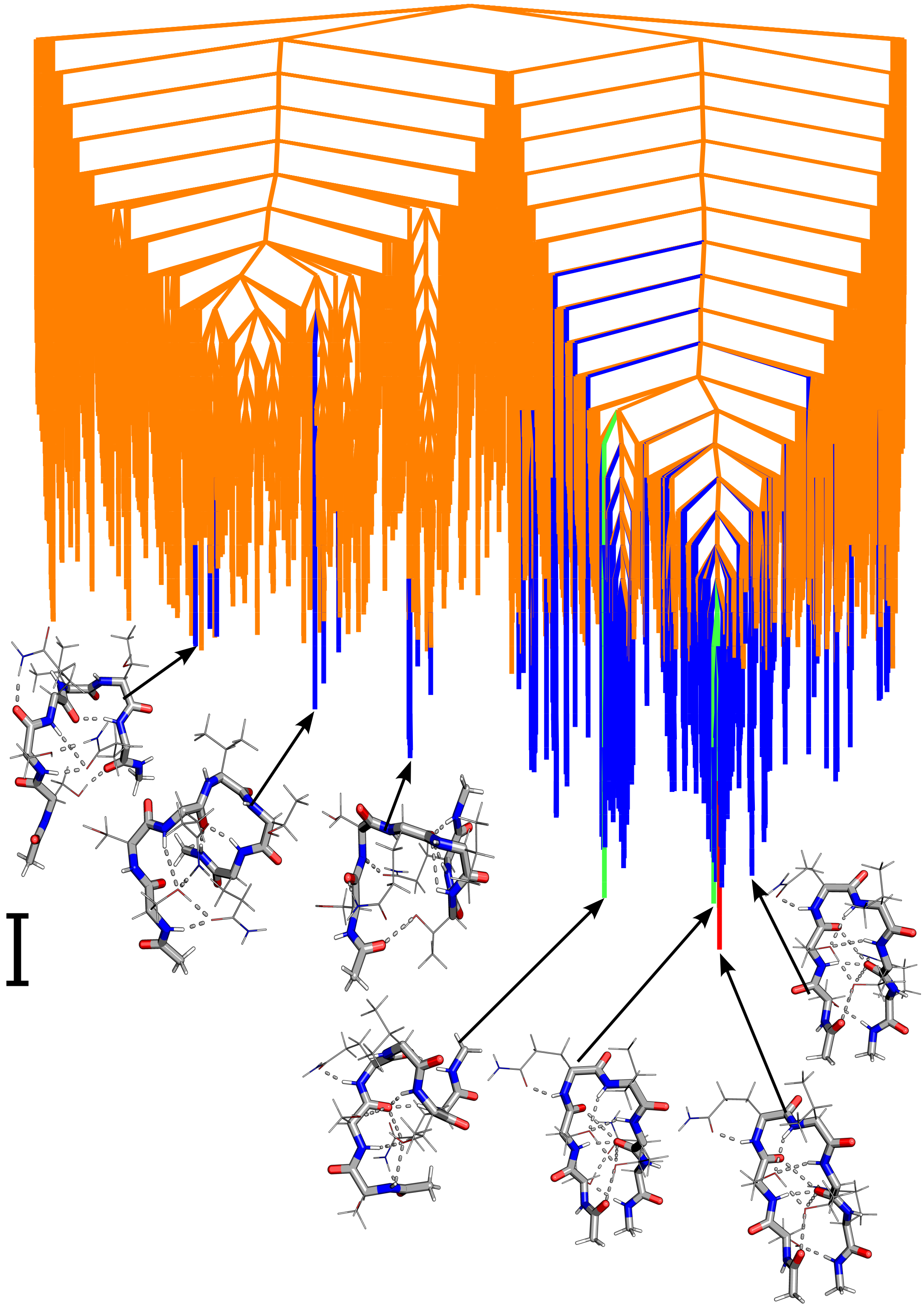

FIG. S21: Disconnectivity graphs for **SSQVTQ** monomer. Local minima representing transition for peaks/inflection points are represented by red to blue (peak 1), green to orange (peak 2), pink to purple (peak 3) and grey to yellow (peak 4). The scalebar represents 1 kcal $mol^{-1}$.

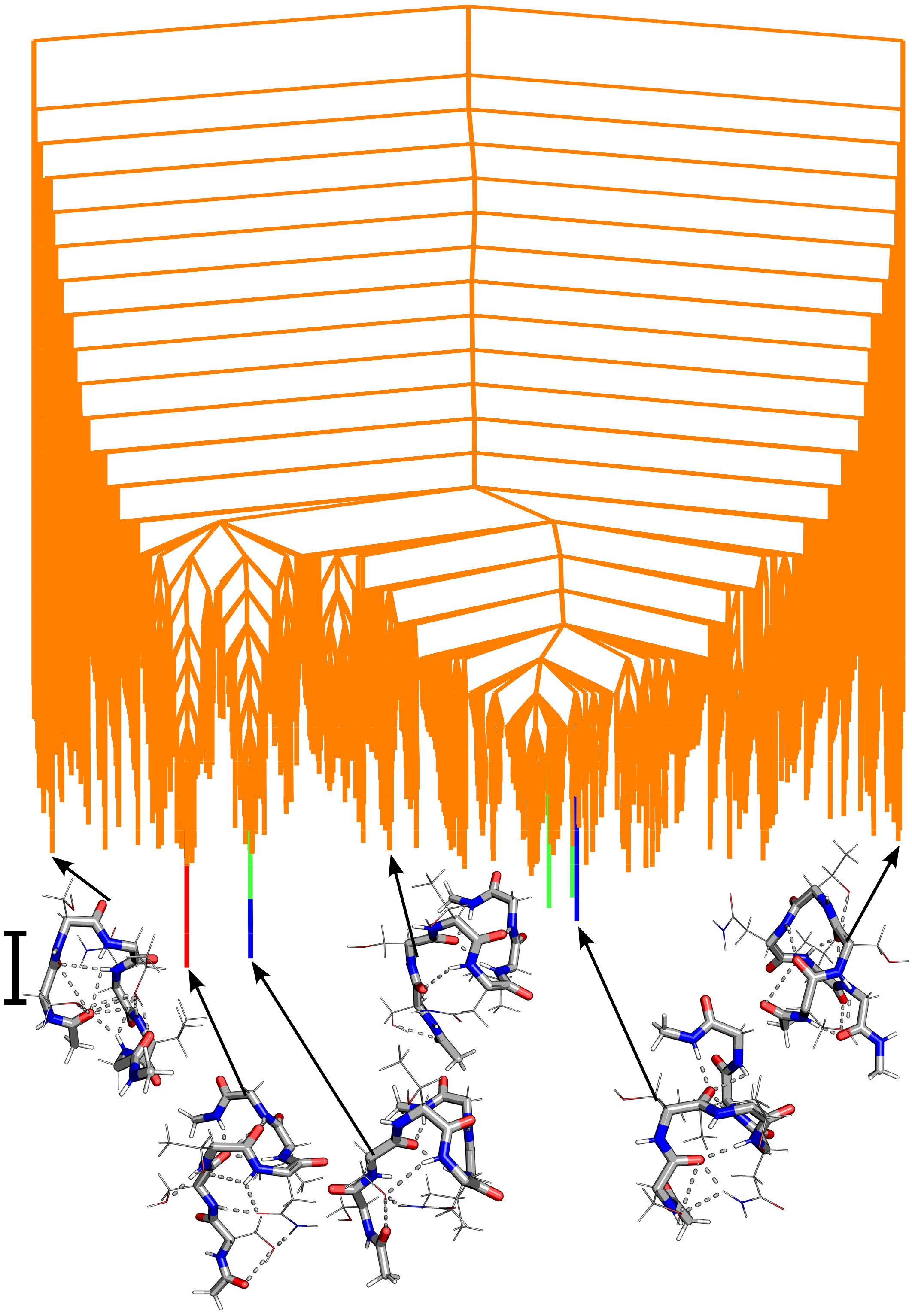

FIG. S22: Disconnectivity graphs for **SSTNVG** monomer. Local minima representing transition for peaks/inflection points are represented by red to blue (peak 1), green to orange (peak 2), pink to purple (peak 3) and grey to yellow (peak 4). The scalebar represents 1 kcal $mol^{-1}$.

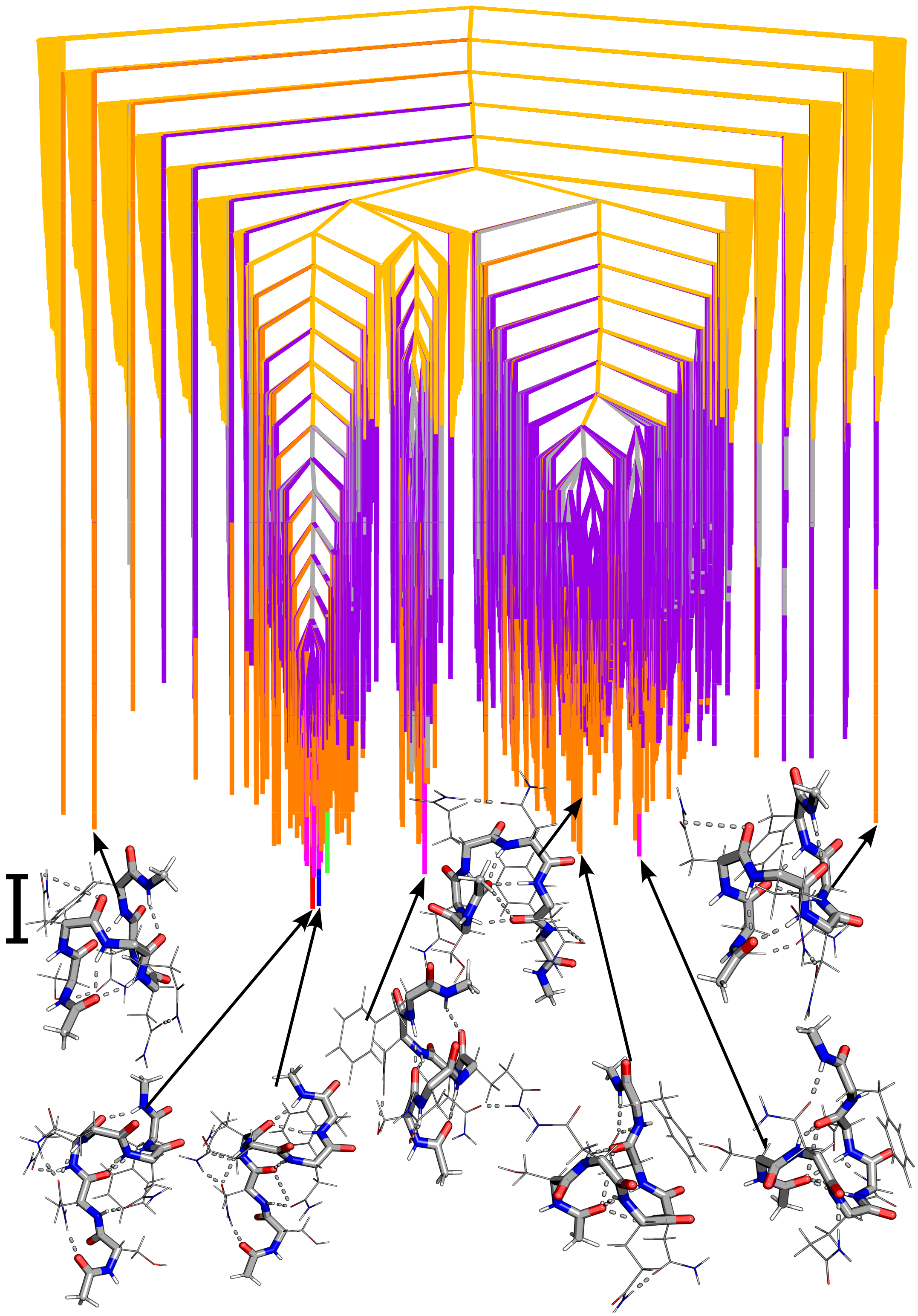

FIG. S23: Disconnectivity graphs for **SNQNNF** monomer. Local minima representing transition for peaks/inflection points are represented by red to blue (peak 1), green to orange (peak 2), pink to purple (peak 3) and grey to yellow (peak 4). The scalebar represents 1 kcal $mol^{-1}$.

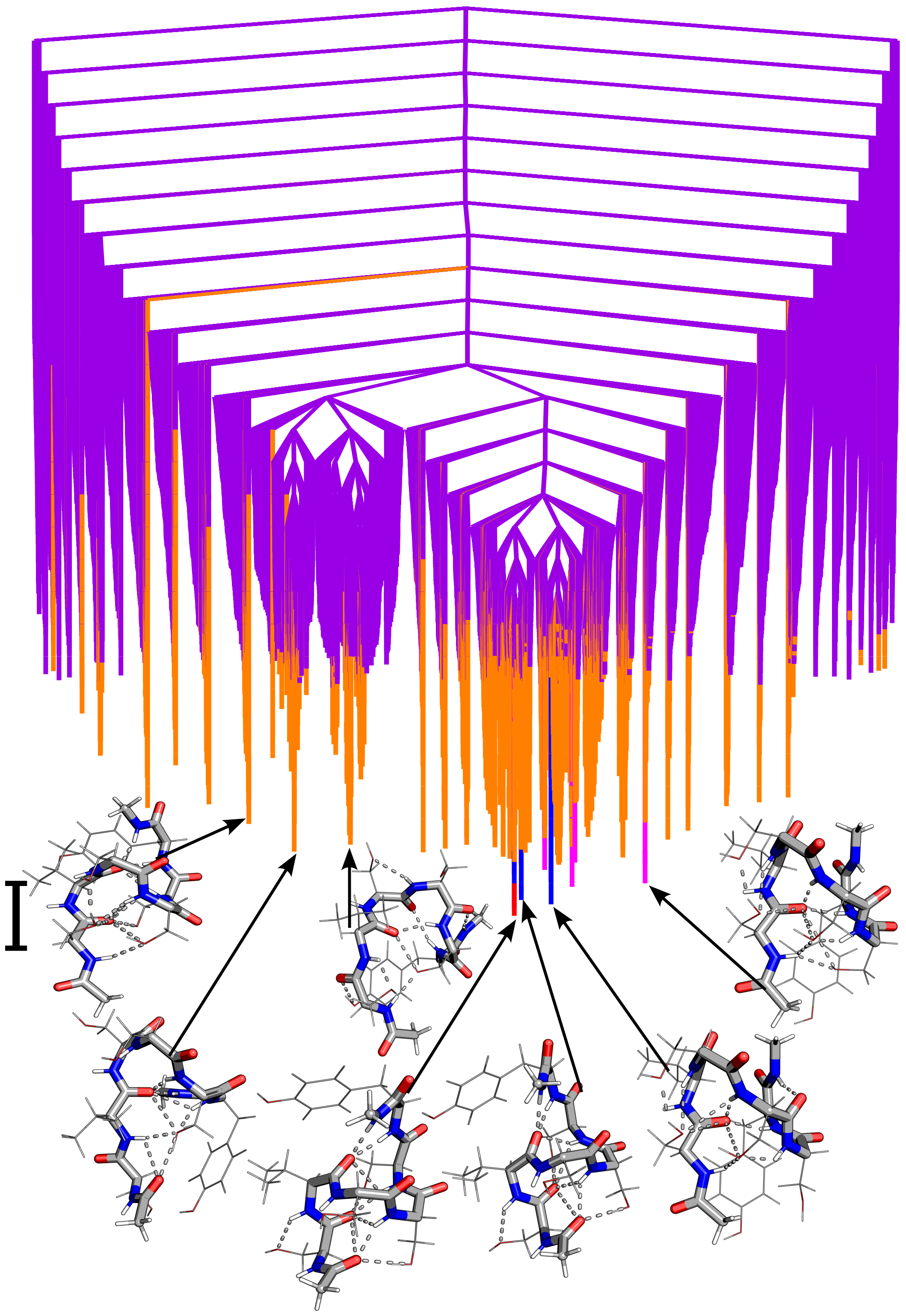

FIG. S24: Disconnectivity graphs for **SVSSSY** monomer. Local minima representing transition for peaks/inflection points are represented by red to blue (peak 1), green to orange (peak 2), pink to purple (peak 3) and grey to yellow (peak 4). The scalebar represents 1 kcal $mol^{-1}$.

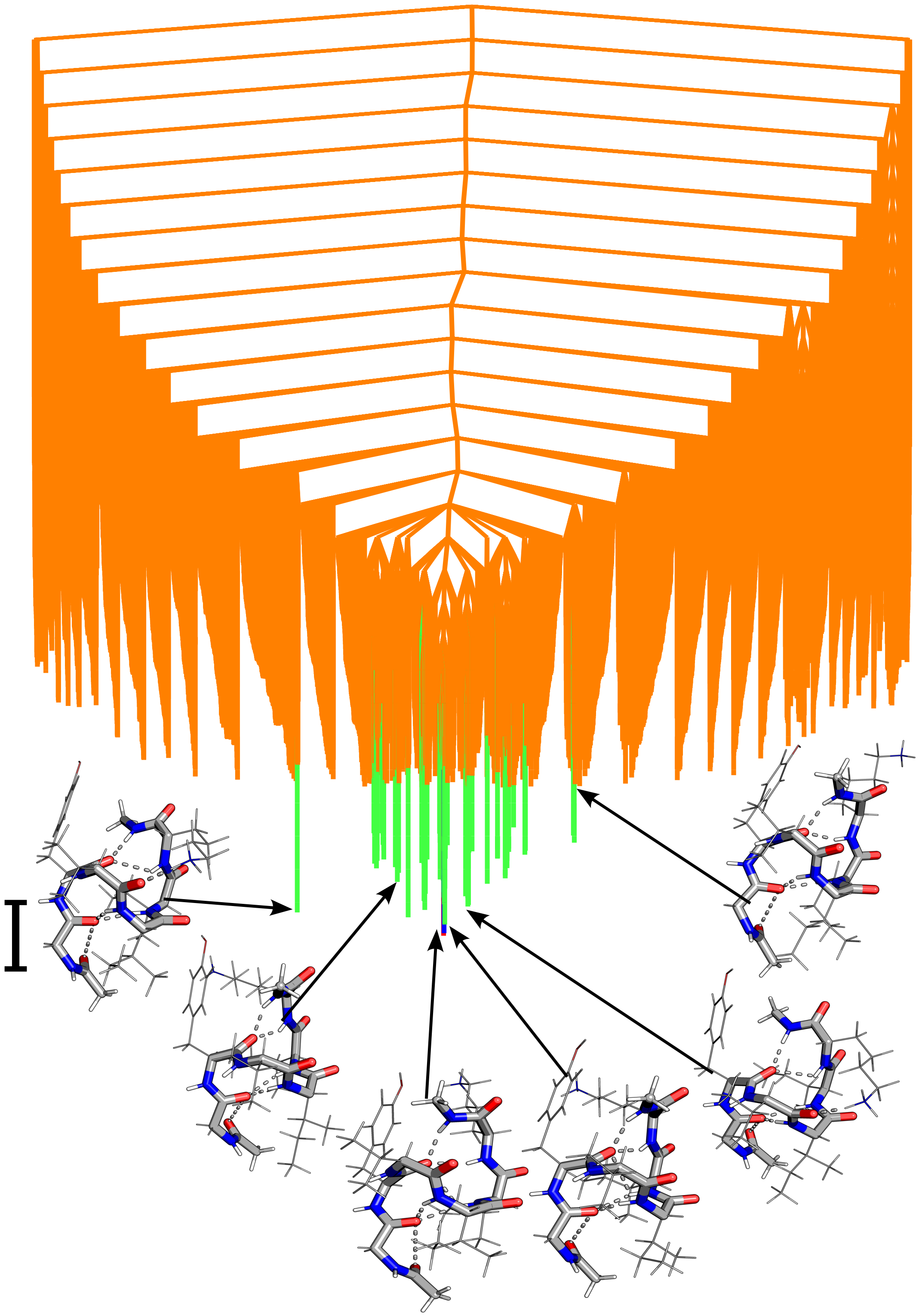

FIG. S25: Disconnectivity graphs for **GYVIIK** monomer. Local minima representing transition for peaks/inflection points are represented by red to blue (peak 1), green to orange (peak 2), pink to purple (peak 3) and grey to yellow (peak 4). The scalebar represents 1 kcal mol$^{-1}$.

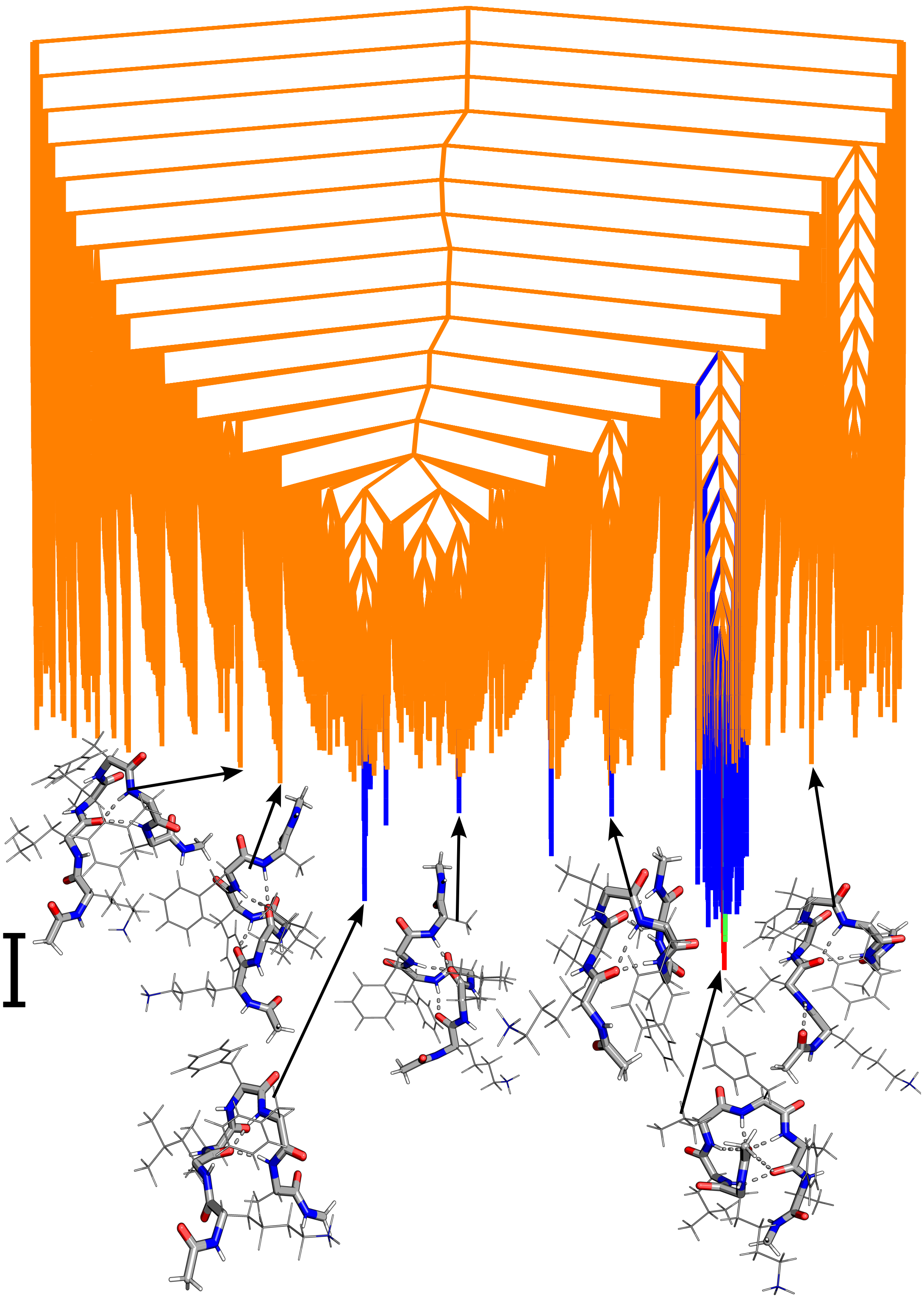

FIG. S26: Disconnectivity graphs for **KLVFFA** monomer. Local minima representing transition for peaks/inflection points are represented by red to blue (peak 1), green to orange (peak 2), pink to purple (peak 3) and grey to yellow (peak 4). The scalebar represents 1 kcal $mol^{-1}$.

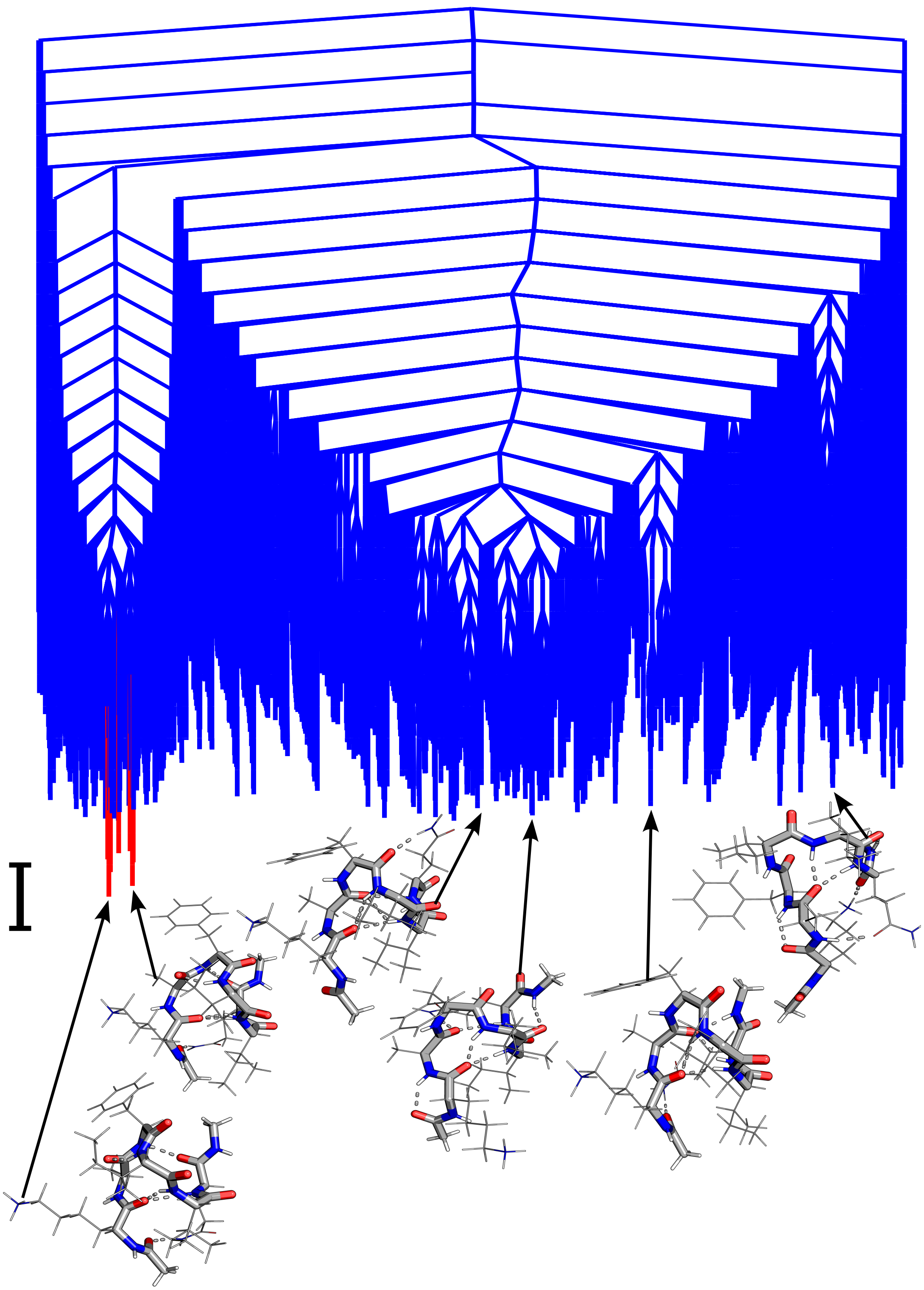

FIG. S27: Disconnectivity graphs for KAFIIQ monomer. Local minima representing transition for peaks/inflection points are represented by red to blue (peak 1), green to orange (peak 2), pink to purple (peak 3) and grey to yellow (peak 4). The scalebar represents 1 kcal $mol^{-1}$.

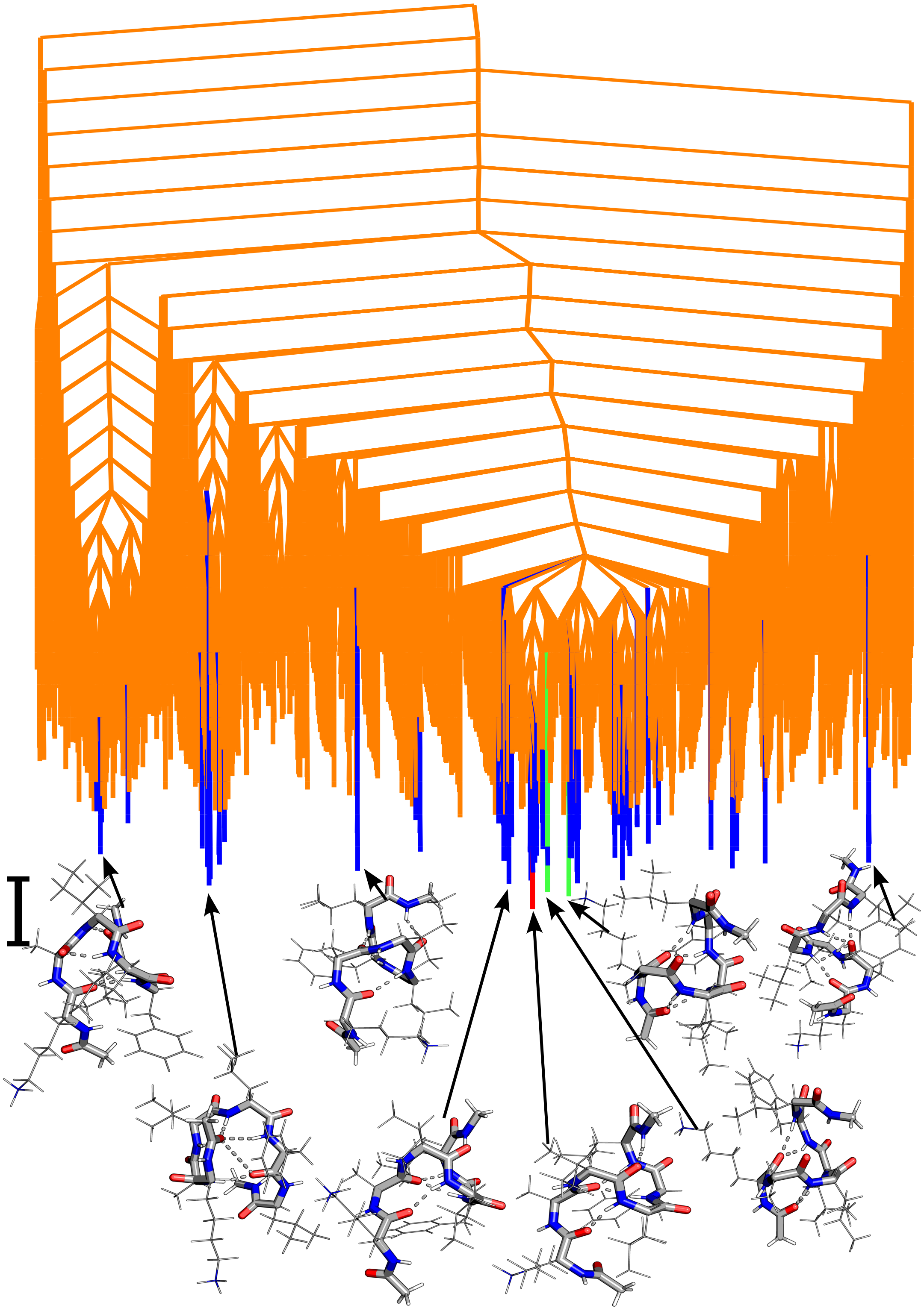

FIG. S28: Disconnectivity graphs for KAILFL monomer. Local minima representing transition for peaks/inflection points are represented by red to blue (peak 1), green to orange (peak 2), pink to purple (peak 3) and grey to yellow (peak 4). The scalebar represents 1 kcal mol$^{-1}$.

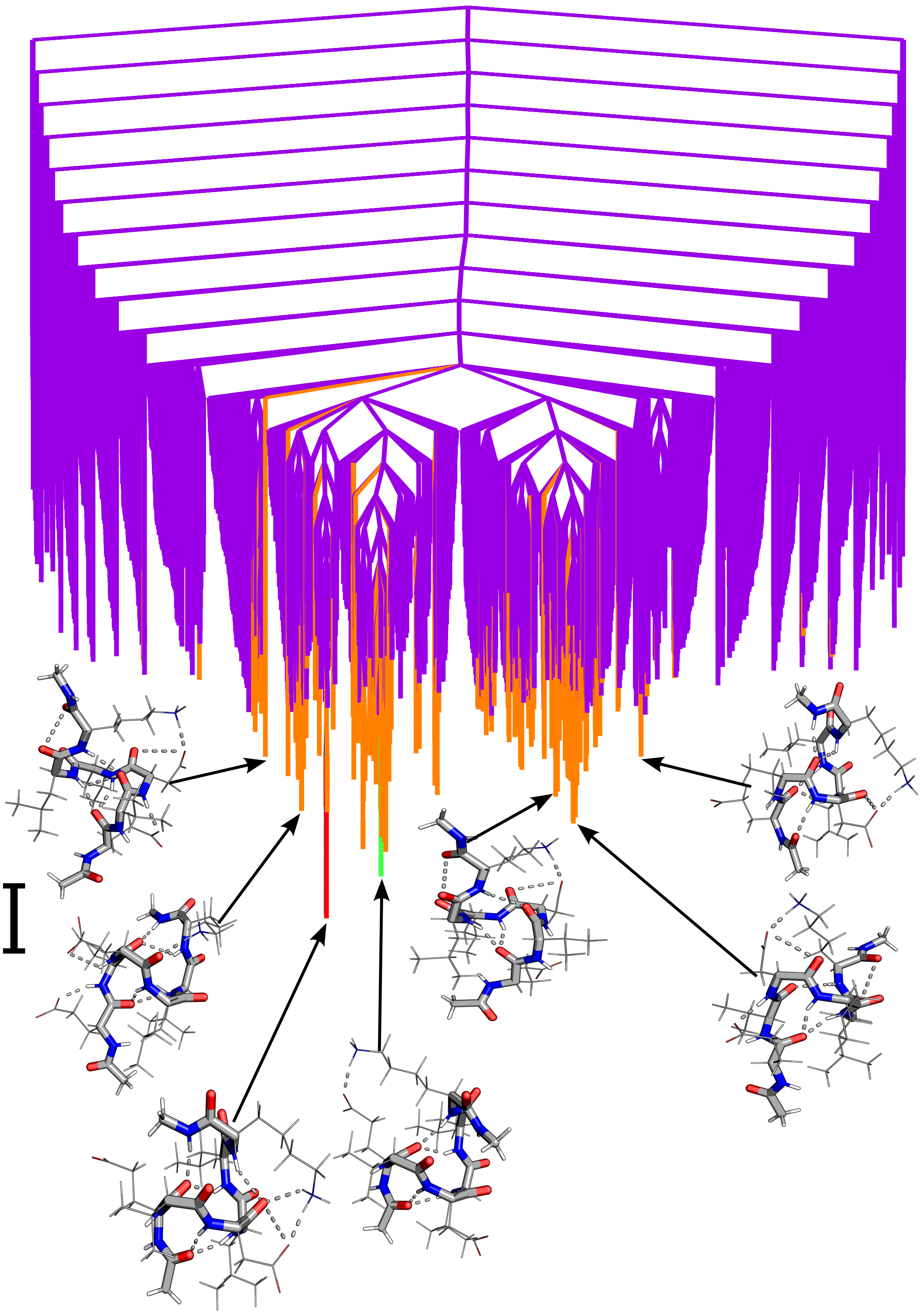

FIG. S29: Disconnectivity graphs for EVDLLK monomer. Local minima representing transition for peaks/inflection points are represented by red to blue (peak 1), green to orange (peak 2), pink to purple (peak 3) and grey to yellow (peak 4). The scalebar represents 1 kcal mol$^{-1}$.

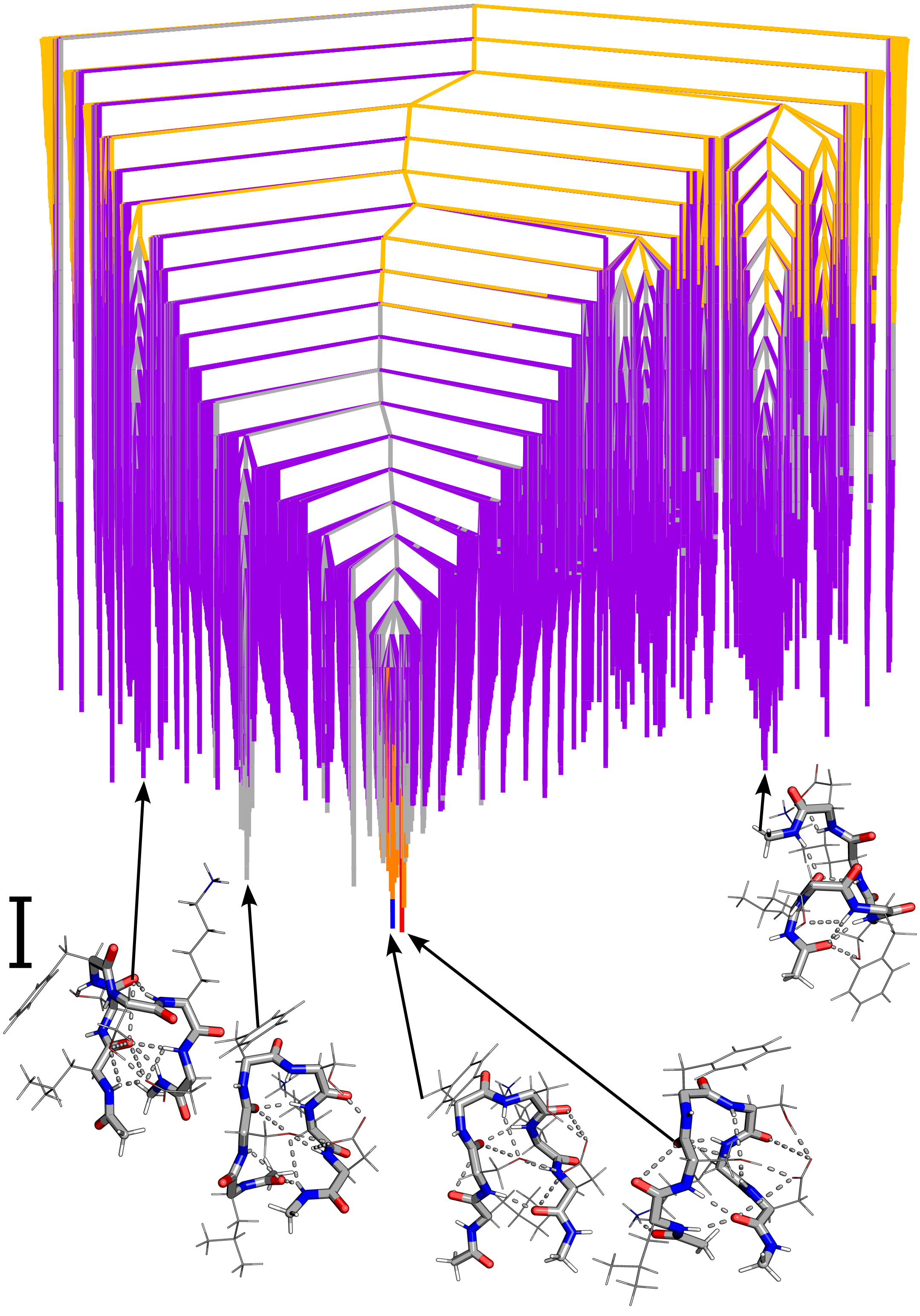

FIG. S30: Disconnectivity graphs for LSFSKD monomer. Local minima representing transition for peaks/inflection points are represented by red to blue (peak 1), green to orange (peak 2), pink to purple (peak 3) and grey to yellow (peak 4). The scalebar represents 1 kcal mol$^{-1}$.

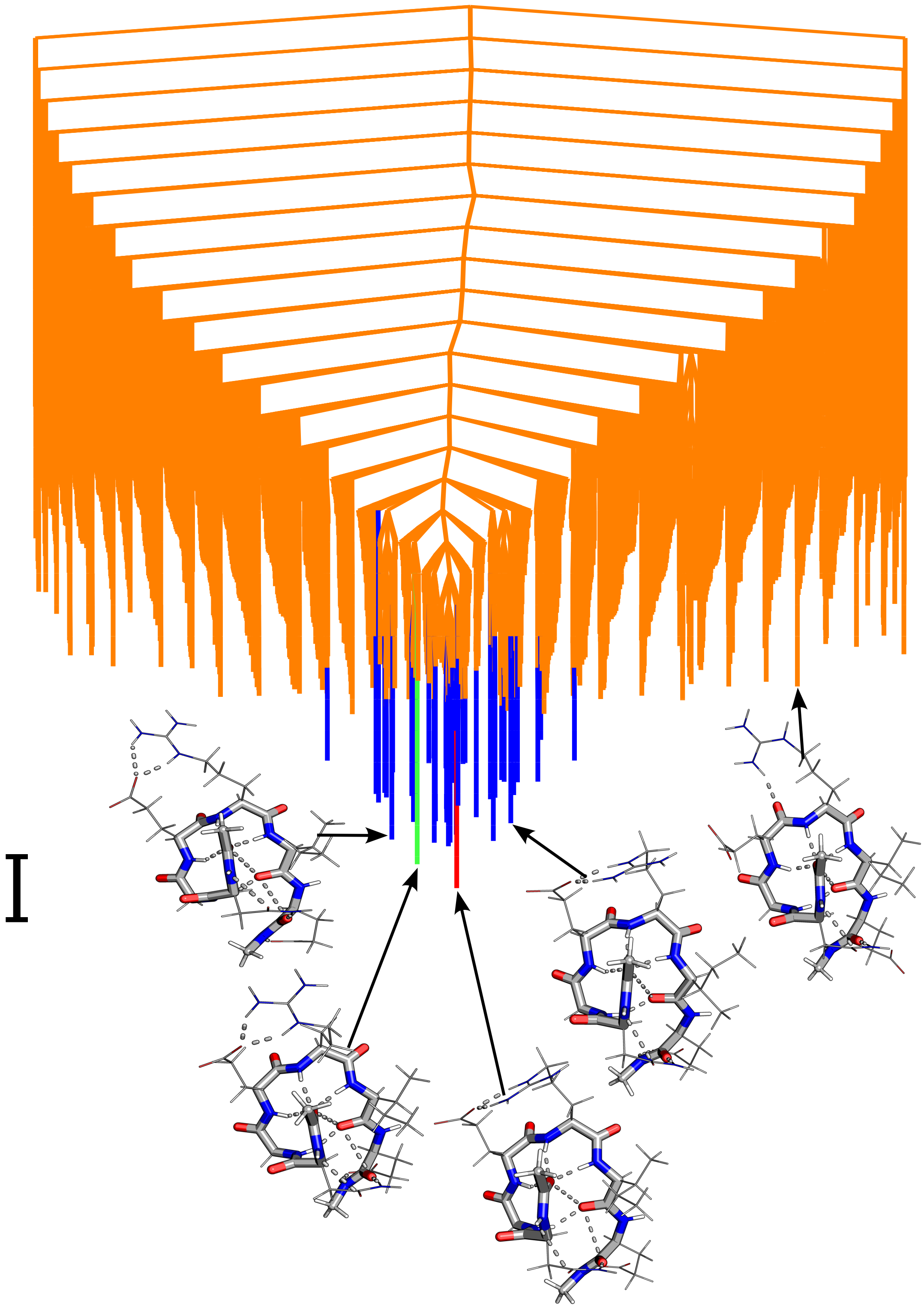

FIG. S31: Disconnectivity graphs for NGERIE monomer. Local minima representing transition for peaks/inflection points are represented by red to blue (peak 1), green to orange (peak 2), pink to purple (peak 3) and grey to yellow (peak 4). The scalebar represents 1 kcal mol$^{-1}$.
